# SampleQC: robust multivariate, multi-celltype, multi-sample quality control for single cell data

**DOI:** 10.1101/2021.08.28.458012

**Authors:** Will Macnair, Mark D. Robinson

## Abstract

Quality control (QC) is a critical component of single-cell RNA-seq (scRNA-seq) processing pipelines. Current approaches to QC implicitly assume that datasets are comprised of one celltype, potentially resulting in biased exclusion of rare celltypes. We introduce `SampleQC`, which robustly fits a Gaussian mixture model across multiple samples, and improves sensitivity and reduces bias compared to current approaches. We show via simulations that `SampleQC` is less susceptible to exclusion of rarer celltypes. We also demonstrate `SampleQC` on a complex real dataset (867k cells over 172 samples). `SampleQC` is general, is implemented in R, and could be applied to other data types.

## Background

Developments in single cell RNA-seq (scRNAseq) technology have enabled experimenters to quantify transcriptional profiles for ever larger and more complex experiments (1; 2; 3). The transformational nature of single cell RNA-seq is its ability to measure transcriptional expression from nanolitre volumes. However, these tiny volumes result in noisy measurement, and the cells themselves can be strongly affected by the process of measurement. Quality control (QC) is therefore a critical element of pre-processing scRNAseq data: experimenters wish to carefully remove ‘bad’ cells, while minimising both loss of healthy cell datapoints and any bias in the types of cells retained (4). Cells can be ‘bad’ due to either technical issues such as droplets with unsuccessful reactions, or biological perturbations such as apoptosis. As the size and complexity of experiments increases, doing this for each sample individually becomes more challenging, and researchers require computational assistance to make the task feasible.

A fundamental issue with methods aiming to exclude poor quality cells is adequately defining ‘poor quality’. This is both challenging in itself and context-dependent. One approach to defining quality is in terms of the quantity of information provided by a cell, i.e. the number of reads or features. We might then seek to exclude all cells with too few reads. However, if there are sufficiently many of these cells, the total information provided could be enough to perform useful tasks (e.g. using ‘pseudobulk’ approaches to identify genes with differential expression between conditions (5; 6)). Similarly, high proportions of mitochondrial reads are associated with stressed and apoptotic cells (7), and excluding cells on the basis of high mitochondrial proportion (e.g. cells with greater than 5% mitochondrial reads in mice, or greater than 10% in human cells; (8)) is a recommended approach to QC (4; 9). However, analysis of healthy tissue shows that the proportion of mitochondrial reads shows strong variation across both species and tissue type (8), suggesting that this fixed-threshold strategy may exclude healthy cells and fail to exclude relatively stressed cells. Users may prefer more sensitive approaches that do not give hard assessments of cell quality, but give them the power to decide which are the unwanted cells.

Based on the various tutorials available, the current industry standards for QC are `scater` (10) and `Seurat` (11), both of which are data-driven rather than based on a fixed threshold. In addition, a probabilistic method `miQC` was recently introduced (12). All three of these are unimodal (`scater` and `Seurat` are also univariate in typical use). `scater` and `Seurat` function by calculating robust measures of centre and scale (such as median and median absolute deviation, or MAD) for each QC metric, then discarding cells that are a specified number of deviations away from the centres. They are typically applied individually to each sample, which we find may introduce bias. Most importantly, where samples are composed of multiple celltypes whose QC metric distributions differ (e.g. they have lower counts, or higher mitochondrial proportions), such filtering will preferentially exclude cells from the celltypes with more extreme values, leading to a bias towards excluding cells from particular celltypes. This was noted by Germain *et al*., who found that stringent filtering on all cells simultaneously led to exclusion of cells that have extreme QC metrics, but not poor quality cells (13). `miQC` is multivariate, assuming that the relationship between features detected and mitochondrial proportion is different in healthy cells and in ‘compromised’ cells. This approach is also unimodal, and may therefore also suffer from a bias against extreme celltypes. Germain *et al*. also found that filtering was necessary, as lenient filtering led to poorer downstream performance (as measured by clustering). Taken together, these findings motivate the need for a QC filtering tool that allows for the presence of multiple celltypes.

As the size of scRNAseq experiments increases, so does their complexity, resulting in ever-higher numbers of conditions and samples measured per experiment (14). This increases the challenge of doing sensitive QC, e.g. by manually deciding a threshold for each sample. In addition, it is important to measure experimental quality at a *sample* and not just a cell level. An experimenter may wish to make comparisons between samples deriving from multiple experimental conditions; this requires establishing that there are sufficient high quality samples of each desired condition.

To address these problems, we have developed `SampleQC`, a computational method designed for QC of large scRNAseq experiments with complex designs. `SampleQC` is implemented as an `R` package based on a robust multivariate Gaussian mixture model (GMM). By fitting a mixture model in which each of the mixture components corresponds to one ‘QC celltype’, `SampleQC` identifies the QC distributions for multiple celltypes simultaneously, removing the bias towards excluding healthy celltypes with extreme QC metric values. We use a fitting procedure that is statistically robust, in the sense that it expects outliers to be present and inference is not distorted by them. By allowing each sample to have its own ‘sample shift’ term, `SampleQC` fits simultaneously across multiple samples, which both increases statistical power and automates an otherwise tedious process. This addresses not only problems with sensitivity of outlier detection, but also provides sample-level QC outputs. Finally, `SampleQC` provides a comprehensive, automatically-generated report on cell and sample quality.

## Results

### SampleQC overview

Examining the distributions of QC metrics across many single-cell experiments, we observed that the values for a given sample would typically follow a multivariate Gaussian mixture distribution, i.e. groups of cells concentrated around distinct mean values, with distances from these mean values following approximately ellipsoid densities (Figure S1). We also observed that multiple samples often showed approximately the same distribution, subject to a sample-level shift in values, i.e. the individual values might be on average higher or lower than other samples, but the shapes of the ellipsoids and the relative distances between their means remained approximately the same (Figure S1). This motivated the following model to describe the metrics: for a cell *i* from sample *j*, which is a member of QC celltype *k* (*k* is not known), *X*_*i*_ ∼ *N* (*α*_0_ + *α*_*j*_ + *β*_*k*_, Σ_*k*_), where *X*_*i*_ is the vector of QC metrics, *α*_0_ is the global mean, *α*_*j*_ is the mean shift of sample *j* relative to the global mean, and (*β*_*k*_, Σ_*k*_) describe mean and variance-covariance of mixture component *k*. Finally, we observed that across large experiments, not all samples would follow the same mixture distribution, but that the samples could be grouped into ‘sample groups’ with similar distributions of QC metrics.

For the metrics to follow a Gaussian mixture model, it is important that the marginal distributions of unimodal data are approximately Gaussian. This is dependent on appropriate transformations of the data, that allow non-Gaussian metrics to be transformed into spaces where they are approximately Gaussian. Applying `log` transformations to library sizes and genes detected is common practice; in addition, we have found that applying the `logit` (inverse logistic) transform to mitochondrial proportions often results in approximately Gaussian distributions (Figure S2). These observations suggest that, after appropriate transformations, a Gaussian mixture model may be a reasonable description of the data.

We assumed that the components of the mixture model represent ‘QC celltypes’, where cells at the centres of the distributions are good quality, and cells that are unusually distant from these centres are cells with extreme QC metric values that should be discarded. In addition, we sought to identify entire samples that were very different from most samples, to flag these to the user for possible exclusion. This motivated an outlier detection algorithm based on the following two steps: (1) identifying groups of samples whose multivariate QC values approximately follow the same distributions; and (2) within each of these, fitting a multivariate GMM to identify cell-level outliers. These steps are outlined in Figure 1.

**Figure 1:**
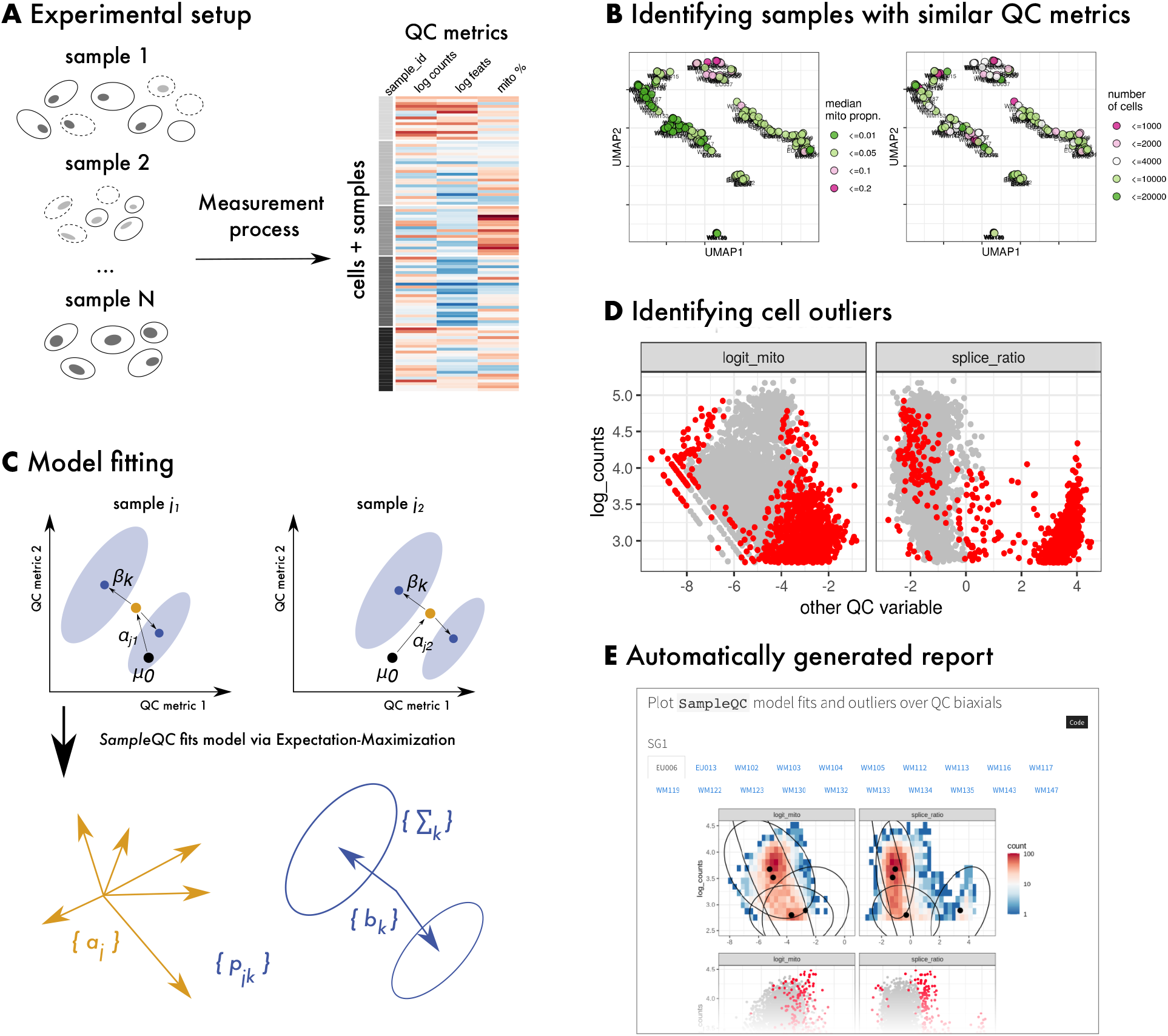
`SampleQC` methodology. **A** Cartoon of cells and samples with varying quality, and their corresponding QC metrics. **B** `SampleQC` estimates distributional distances between samples, and uses this to identify groups of samples. **C** `SampleQC` fits a robust GMM to each group of samples. **D** Low likelihood is used to exclude poor quality cells. **E** `SampleQC` generates report of outliers and overall sample quality.

To identify groups of samples with similar distributions, `SampleQC` uses a non-parametric statistical measure of dissimilarity between multivariate samples, maximum mean discrepancy (MMD: (15)), to estimate ‘distances’ between samples. MMD takes two samples as input, which must have the same dimensions but may have different numbers of observations, and calculates a measure of distance/dissimilarity that summarizes differences in means, (co)variances and higher moments. We then use pairwise distances to construct a nearest neighbour graph that allows the identification of ‘sample groups’, i.e. groups of samples following approximately the same QC metric distributions. We use Louvain clustering to identify the groups (16). The distances are also used as input to dimensionality reduction techniques, such as multidimensional scaling (MDS: (17)) and UMAP (18). This gives low-dimensional embeddings where each point corresponds to a sample, and permit users to check for batch effects or other issues with the data. For example, users can check the extent to which samples from the same batch are clustered together, potentially indicating sample quality problems with a batch.

The second step identifies cell outliers, i.e. cells with very different QC metric values to most other cells. Additionally, this step checks for outlier samples by estimating the mean shift in QC statistics for each sample. After `SampleQC` has identified sample groups, the user then determines how many components are present in each sample group by inspecting the QC distributions within each group, i.e. how many components the GMM should have. `SampleQC` then fits a robust GMM to the cells in each sample group.

Our GMM makes two main assumptions: (1) the configuration of QC celltypes (i.e. their covariance matrices and relative means) is constant across the samples in a sample group; and (2) each sample has a uniform shift in (transformed) QC metrics. Our justification for these assumptions is the observations on distributions of QC metrics discussed above, and the empirical performance of the method, although reasonable biological arguments can also be made for them. The model allows the proportions of QC celltypes (i.e. mixture components) to vary by sample; details of the mathematical model are discussed in Methods (see `SampleQC` model). This model structure, in which the same QC celltypes are assumed to be present in every sample, has benefits when identifying outlier cells. The model learns the parameters for the QC celltype components across the *whole* sample group, so that even when a sample has only a small proportion of a given QC celltype, these cells are not regarded as outliers. These cells may in fact be of particular interest, being from a celltype with reduced proportion relative to other samples.

We fit this model to identify poor quality cells, namely those that are outliers with respect to the distribution of the bulk of cells. While the presence of outliers is known to affect the estimation of statistics, especially higher order moments such as covariance (19), accurate estimation of the covariance matrices is essential precisely for identifying those outliers. We therefore use a *robust* estimation of the mean and covariance matrices (20): robust estimators (like the MAD) give good but slightly less than optimal results when the data do follow an assumed distribution, but are also guaranteed to give good results when the data is contaminated and MLEs may fail (19). We explicitly assume that outliers may be present, making this approach necessary.

Once we have fit models to each sample group, `SampleQC` is able to describe all cells in the included samples in terms of mixture components, which we assume to represent good quality cells. We can then identify outlier cells by calculating a statistical measure of distance from the centre of each QC celltype (Mahalanobis distance, which indicates how unlikely a particular observation is under a *χ*^2^ distribution), and exclude cells that are unlikely to be a member of any component.

### Simulating QC data for a large experiment

With single cell data, there is no ground truth regarding which cells or samples are poor quality, making assessment of the performance of QC methods challenging. To evaluate `SampleQC` against other approaches, we therefore developed a simulation framework that replicates the distributions of QC metrics observed in biological datasets (pseudocode provided in Methods; see Simulations). The framework and parameters are derived from large and complex experimental designs, including outliers, and are based on a hierarchical model.

At the experiment level, we make the following assumptions: (1) the experiment is composed of a number of sample groups with similar QC distributions; (2) there are a small number of ‘QC celltypes’ present in the experiment. By ‘QC celltype’, we mean a cluster of cells that is distinct within ‘QC space’, i.e. the selected QC metrics. We expect that multiple true celltypes, i.e. clusters that are distinct in terms of expression, may have the same distribution in QC terms. These QC celltypes are assumed to be common across all groups of samples, with a different subset present in each sample group. For each group, we randomly sample samples and cells, and which celltypes are present. We also sample the means and covariances for the QC celltypes, {*β*_*k*_, Σ_*k*_}.

Within each sample group, we then model outliers and sample shifts at the level of samples. We assume that for each sample *j*, there is a uniform shift in the (transformed) QC metrics, *α*_*j*_. This affects all cells in a sample equally. We also assume that each sample has different proportions of the QC celltypes, sampled from a Dirichlet distribution. Each sample has a different proportion of outliers, modeled as a beta distribution, which gives all samples from the same group outlier proportions that vary around a common mean value. Outliers are assumed to differ from ‘good’ cells by having lower non-mitochondrial reads and numbers of features, and correspondingly higher mitochondrial proportions; we model this by a sample-level parameter corresponding to the proportion of non-mitochondrial reads lost (modelled as a logistic normal).

For every cell, we have annotations of sample group (*g*), sample (*j*) and QC celltype (*k*), with corresponding values of global mean (*µ*_*g*_), sample shift (*α*_*j*_) and QC celltype mean and covariance (*β*_*k*_ and Σ_*k*_). We draw a healthy QC metric vector for a given cell via a multivariate normal with mean *µ*_*g*_ + *α*_*j*_ + *β*_*k*_, and covariance Σ_*k*_. We then determine whether this cell is an outlier via a Bernoulli distribution, and if it is, we perturb this value by downsampling the non-mitochondrial reads and the QC metrics are updated accordingly.

This approach follows the model assumed by the GMM (`SampleQC` model), with two additional points to make the simulations more challenging. Firstly, the GMM does not include any outliers. To model these, we assume that each sample has a given proportion of outlier cells, selected at random. For each of these cells, the non-mitochondrial reads and the number of features are decreased by a given proportion, according to a beta distribution. This results in both a reduction in total counts and features, and an increase in mitochondrial proportion, consistent with outliers observed in real data. Secondly, we assume that the relative positions of the components, {*β*_*k*_}, are not identical in each sample, and vary by a small amount, *δ*_*jk*_.

This results in simulated QC metrics for tens of thousands of cells across tens of samples, where the distributions of these metrics vary coherently across different groups of samples. The simulations provide two matrices of QC metrics: one that mimics the QC values we might see from experiments, and one for the QC values of the cells ‘before’ they became low quality (such a matrix would never be observed in real life).

### Benchmarking SampleQC against scater and miQC on simulated data

We generated simulated data for an experiment with 100,000 cells across multiple samples, resulting in 64 samples with a median of 1,300 cells per sample, including outliers. We then ran `SampleQC`, `scater` and `miQC` to compare performance: both in terms of excluding outliers, and retaining ‘good’ cells. Inspecting the distributions of the outliers identified by the different methods, we observe that where multiple QC celltypes are present, `SampleQC` identifies outliers correctly, while both `scater` and `miQC` may preferentially exclude cells from one QC celltype as outliers (Figure 2). Where only one QC celltype is present, `SampleQC`, `scater` and `miQC` show similar results (Figure S3), as expected.

**Figure 2:**
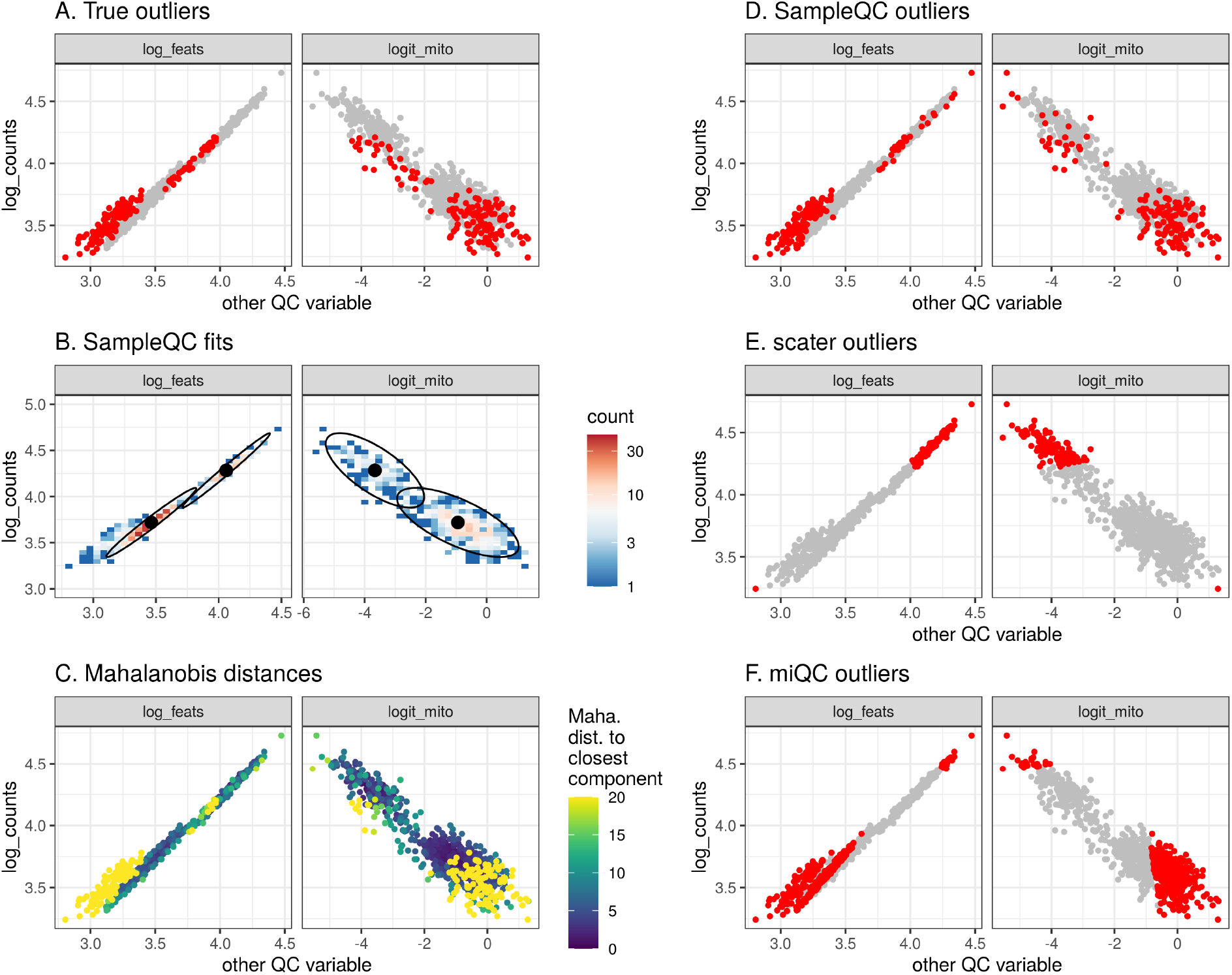
QC method comparisons on multimodal simulated data. **A** Biaxial plot of QC metrics for one simulated sample. Each point represents the QC metrics for one cell, with (known) simulated outliers in red and non-outliers in grey. **B** Multivariate density plot of data in (A), annotated with models fit by `SampleQC`. The distinct clusters here are examples of what we term ‘QC celltypes’, i.e. cells with similar distributions in their QC metrics, although their gene expression distributions may be distinct. **C** Mahalanobis distances to nearest cluster under the fit `SampleQC` model, with large values indicating low likelihood under multivariate Gaussian mixture model. **D, E, F** Outliers detected by `SampleQC`, `scater` and `miQC` respectively. These indicate the different assumptions on outliers made by the different methods: `SampleQC` calls droplets in low-density regions of the QC metric space as outliers; `scater` calls droplets with extreme values on at least one metric as outliers, in effect drawing a box around the central mode of the data; `miQC` identifies droplets with high mitochondrial proportion relative to the number of features observed, which may correspond to a good quality celltype with high mitochondrial proportion.

When we assess performance in identifying good cells across the entire simulated dataset, we find that `SampleQC` has equal or better performance than both `scater` and `miQC` for both precision and recall (Figure 3A, Figure S4). The relative performance on individual samples is affected by how many celltypes are present in the sample. Where the data is multimodal, the multimodal model in `SampleQC` gives it equal or better performance than `scater` and `miQC` for both precision and recall (Figure S4, Figure S5). Where the data is unimodal (i.e. has one mixture component), `scater` and `SampleQC` show similar performance (Figure 3B). In this case, performance of `miQC` is slightly worse, potentially because it assumes that a reasonable quantity of ‘compromised’ cells must be present; when there are only few such cells, this may result in ‘good’ cells reported as outliers.

**Figure 3:**
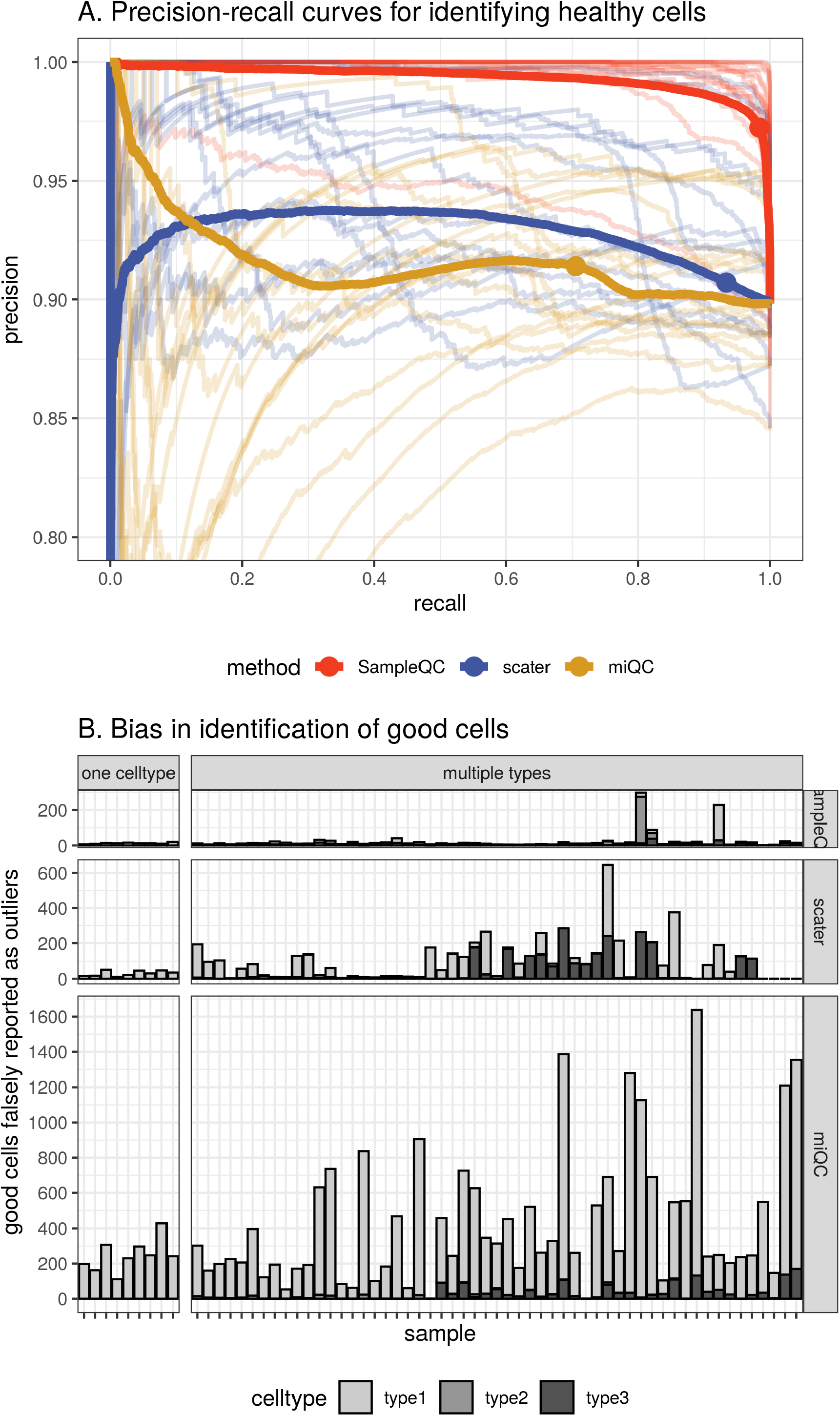
Performance comparison of `SampleQC`, `scater` and `miQC` on simulated data. Results from a simulated experiment of 100k cells with 3 QC celltypes. **A** Precision-recall curve of identification of ‘good’ cells. Solid lines show curve calculated over all cells. Transparent lines show curves for individual samples. Dots show default exclusion thresholds for each method (*<* 1% Chi-squared likelihood under fitted model for `SampleQC`; *>* 2.5 MADs for `scater`; *>* 0.75 posterior probability of compromised cell for `miQC`). **B** Bias in number of cells reported by sample, samples split by whether one or multiple QC celltypes present in the sample (i.e. whether samples is unior multimodal). *y*-axis is bias in reporting of good cells.

Importantly, `SampleQC` also shows better performance than the compared methods in terms of celltype bias. As above, when one QC celltype is present in a sample, `SampleQC` and `scater` have very similar biases in excluding good cells at default settings. However, when multiple QC celltypes are present in a sample, `SampleQC` excludes similar numbers of good cells, and shows low bias towards any particular QC celltype. In this setting, the good cells mistakenly excluded by both `miQC` and `scater` are almost entirely comprised of one celltype, while `SampleQC`’s more flexible model better preserves rare QC celltype proportions. We note that these results are dependent on the choices made in developing the simulation, and that alternative reasonable choices may give different results.

We performed the same comparisons for simulations where the data is t-distributed, and found similar differences in performance between the methods (see dashed lines in Figure S4).

### Number of QC clusters

Determining the appropriate number of distinct clusters in a dataset is a well-studied problem in statistics that remains challenging (21). To avoid this problem, `SampleQC` requires users to specify the number of mixture components for each group of samples. We simulated experiments to test the sensitivity of our results to mismatches between the selected and true *k*. We simulated data with different true numbers of QC celltypes, varying over 1, 2 and 3 QC celltypes (our observation over many studies has been that many samples have between 1 and 4 mixture components). For each value of `k_true`, we simulated data for 20,000 cells divided into samples, with an average of 2000 cells per sample. We then fit `SampleQC` to each dataset, with the assumed number of clusters `k_fit` varying between 1 and 4. We found that a mismatch between `k_true` and `k_fit` could result in a reduction in performance (while `SampleQC` performed well when `k_fit` was correctly specified) (Figure S7). However, for multiple of the combinations where `k_fit` was misspecified, `SampleQC` was not able to fit; this provides some protection in itself. These results indicate that review of the diagnostic plots rendered by `SampleQC` is an important part of the QC process, allowing users to determine plausible values of *k* from the empirical distributions.

### Applying SampleQC to complex biological datasets

We applied `SampleQC` to a large single-nuclei RNA-seq dataset comprising 867k cells over 172 samples (see Data availability), taken from human brain tissue, including samples from patients with neurodegenerative conditions. This illustrates an intended use case for `SampleQC`: a dataset with sufficiently many samples to make it challenging to adjust outlier thresholds for each sample individually, and where complex batch effects could be present. Here, we use log library size, logistic transform of mitochondrial proportion; and log2(spliced reads / unspliced reads) as QC metrics. The motivation for using log splice ratio in single nuclei data comes from the difference in abundance of unspliced reads between the nucleus and the cytoplasm (22). In principle, unspliced reads should only be present in the nucleus; high ratios of spliced to unspliced read counts therefore allow us to identify nuclei that were not successfully stripped of their cytoplasm, and are likely to be contaminated. In addition, this demonstrates the flexibility of `SampleQC`, which only assumes that multiple QC celltypes are present, and does not require specific metrics to be used.

`SampleQC` first identifies samples with similar QC distributions, by calculating pairwise MMD distances between the samples. The principle reason for this is to allow `SampleQC` to fit separate models to each group of individual samples. Low dimensional projections of the resulting dissimilarities are shown in Figure 4 (UMAP) and Figure S8 (MDS), showing the groups identified by `SampleQC` (Figure 4A), allowing investigators to assess whether sample source, sequencing run or other covariates might have influenced sample quality (Figure 4B). Investigators can also check whether there are groups of samples that should be excluded entirely due to poor quality.

**Figure 4:**
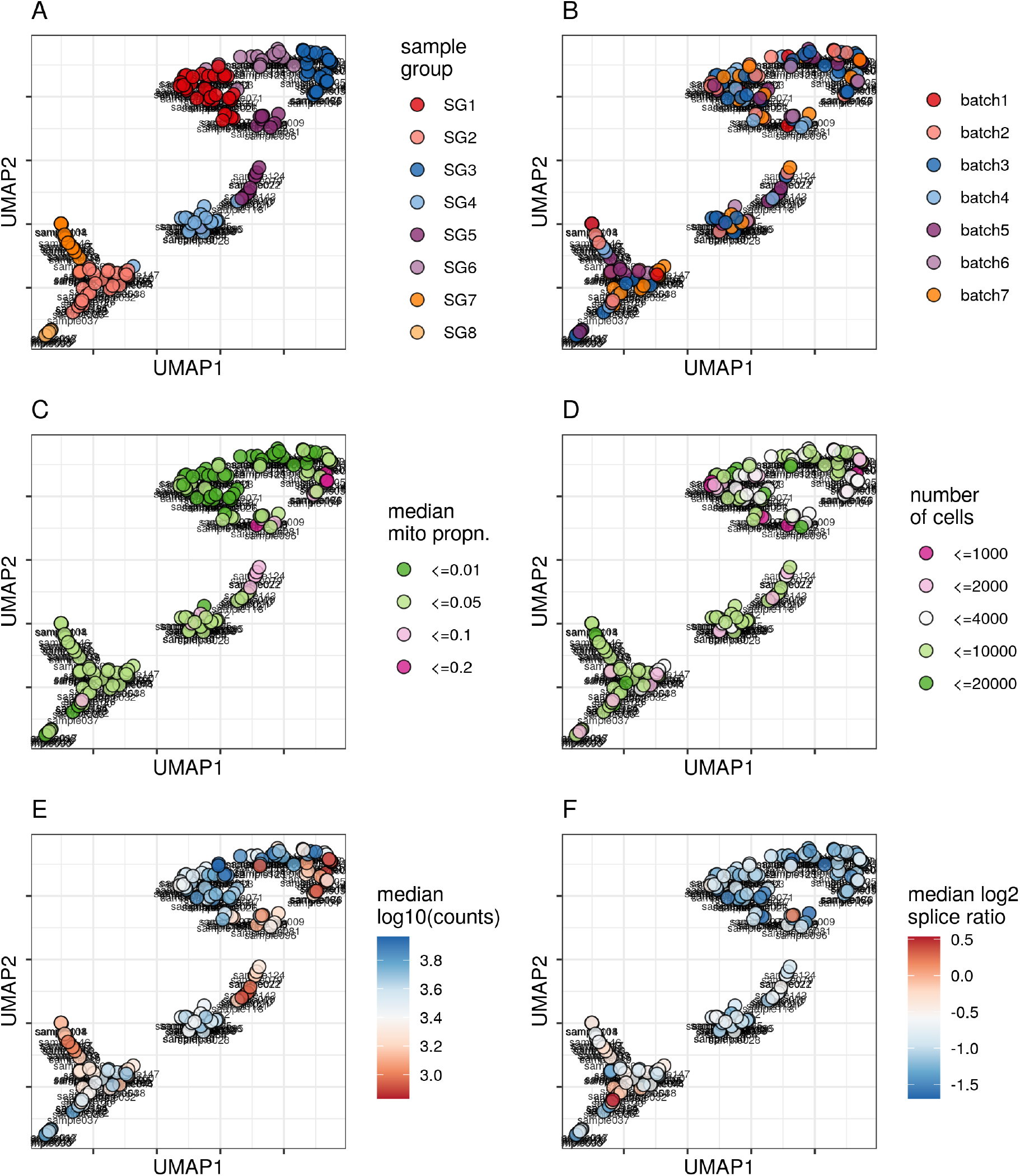
Sample-to-sample distance embedding (UMAP). `UMAP` embedding, based on sample-sample similarities calculated with MMD, resulting in samples with similar QC metrics being plotted close to each other. **A** Sample group, i.e. clusters of samples with similar QC metrics distributions, as estimated by `SampleQC`. Remaining plots show sample metadata and statistical summaries: **B** Sequencing batch **C** Mitochondrial proportion (categorized) **D** Number of cells (categorized) **E** Median counts per cell **F** Median log2 splice ratio per cell.

Applying `SampleQC` to this dataset shows similar results to those for the simulation: where a cell population is present in the whole dataset but only makes up a small proportion of a sample, `SampleQC` is better able to preserve these cells than `scater` and `miQC` (Figure 5). The Mahalanobis distances calculated under the `SampleQC` model also follow the expected distribution, i.e. most points follow a Chi-squared distribution with degrees of freedom equal to the number of dimensions, plus an outlier component with larger distances (Figure S9).

**Figure 5:**
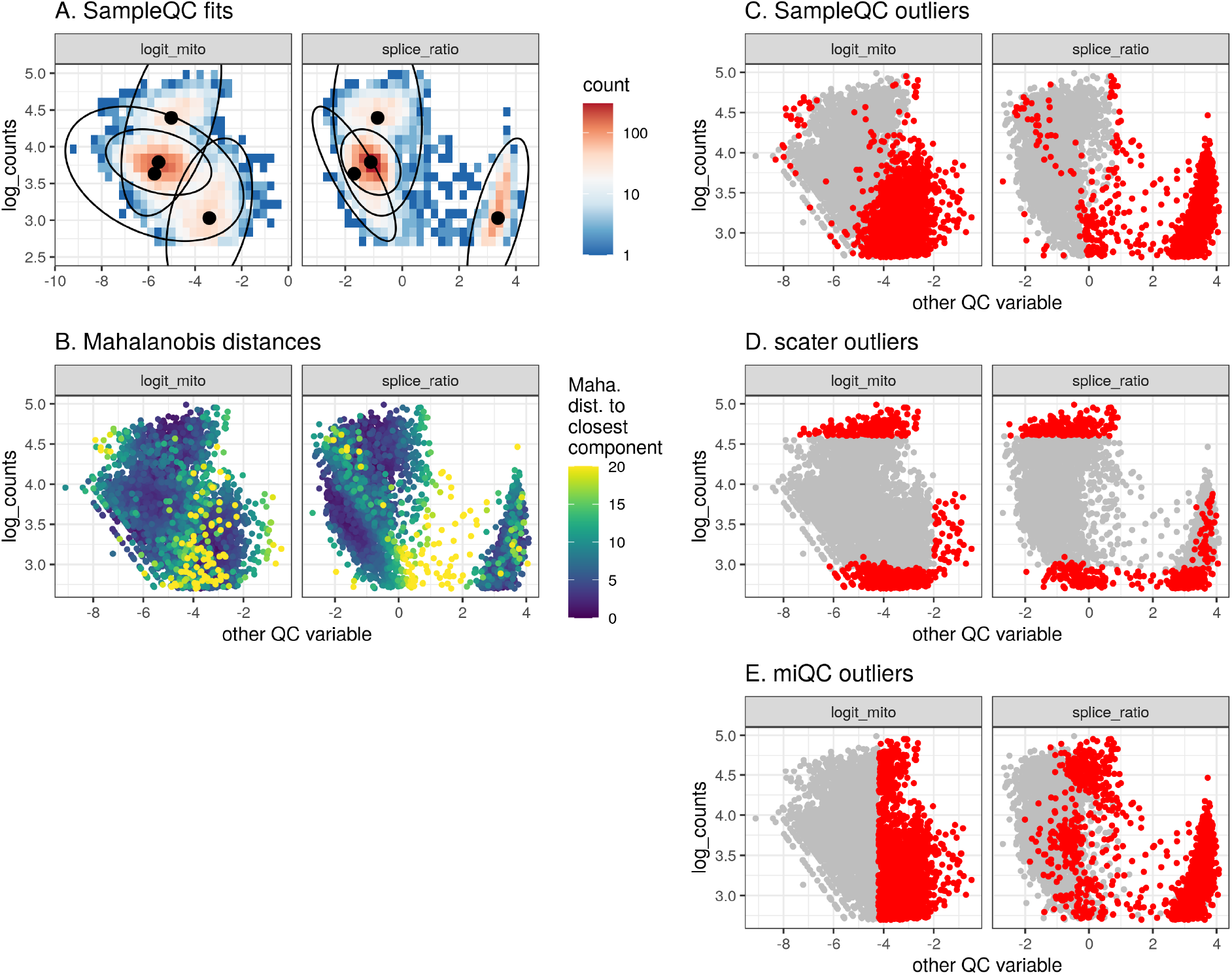
QC method comparisons on complex biological data. **A** Biaxial empirical density plot of QC metrics for single nuclei data, showing one selected sample, annotated with models fit by `SampleQC`. Colour indicates density of cells with the same QC metrics. **B** Mahalanobis distances to nearest cluster under fit `SampleQC` model, with large values indicating low likelihood under multivariate Gaussian mixture model. **C, D, E** Outliers detected by `SampleQC`, `scater` and `miQC` respectively. `SampleQC` outliers include user specification that QC celltypes with mean splice ratio *>* 3 should be excluded.

An alternative approach to QC for single cell data is to apply only a minimal filter on QC metrics (to remove droplets that contain very little information and / or contain only lysed cells), cluster the cells as is normal for single cell, then remove clusters that show poor QC metrics. This approach works for some datasets, however it may become difficult to apply in more challenging samples. Applying this approach to single cell data from ovarian cancer tissue (12), we found that while some of the clusters should clearly be excluded, other clusters showed a wide range of QC metrics (Figure S11). For example, clusters K06, K07 and K08 all include substantial numbers of cells with mitochondrial percentages both above 25% and below 10%, showing that this approach to excluding poor quality cells may be more difficult in samples not comprising healthy cells.

Although `SampleQC` is a more complex model than other approaches to QC, this does not substantially decrease speed. As an illustration, we timed `SampleQC` on the simulated dataset in Figure 3, comprising 100k cells over 64 samples. The three steps of `SampleQC` took the following times: 44s to calculate pairwise MMDs; 70s to fit the `SampleQC` models; and 104s to render the report.

The methodology underlying `SampleQC` is general, and not specific to scRNAseq. The example above demonstrates its application to single nuclei RNAseq, and in principle it can identify outliers in any dataset comprising a small set of mixture components described by a small set of variables. This includes other single cell technologies, such as CITE-seq, which is similar to scRNAseq but also quantifies protein expression via oligo-tagged antibodies (23), and scATACseq, which quantifies chromatin accessibility in single cells (24). To demonstrate this, we ran `SampleQC` on CITE-seq data from PBMCs (25), using similar QC metrics to scRNAseq: log RNA counts; log RNA features; logistic transform of mitochondrial proportion; log ADT (protein) counts; log ADT features. The models fit by `SampleQC` successfully converged to the modes of the data, and could be used for outlier detection (Figure S10).

## Discussion

We introduced `SampleQC`, an approach for quality control of scRNAseq experiments that improves on the current industry standard, and is designed for large and complex experimental designs. We show that our model simultaneously allows flexibility and sensitivity, and reduces bias. Grouping samples by similarity of QC metric distributions allows a different model to be used for each sample group, which then increases sensitivity by borrowing strength across samples.

One of the main challenges in this work is the lack of ground truth to evaluate methods. For this reason, previous QC filtering methods have not performed comprehensive benchmarks. Meanwhile, most datasets are annotated to celltypes after filtering and therefore not annotated as good/bad for such an assessment. And while filtering is necessary at some level, it is also not always straightforward to assign a reason for a cell’s exclusion. For example, many of the excluded cells according to `SampleQC` are in low-density regions, but it could be they are not too extreme in each univariate QC metric, but reside in a low-density region in a multivariate sense. Furthermore, none of the tools available for QC filtering, including `SampleQC`, perform rigorous validation that low-quality cells are truly removed.

Our method is based on empirical observations of real data, and is motivated by known biological mechanisms. Many single cell techniques assume that scRNAseq data comprises multiple celltypes that are distinct in terms of gene expression, however this is not reflected in current approaches to QC. Metrics used for quality control are functions of the full expression vector of counts. When multiple celltypes are present in the full expression data, our observations over multiple datasets are that at least some separation into ‘QC celltypes’ is typically preserved in the restricted ‘QC space’. Our motivations regarding outliers, i.e. that they form a small proportion of the dataset with lower read counts and higher mitochondrial proportions, are also consistent with this observation. `SampleQC` allows users to choose any metric for input, and if the selected metrics approximately follow a GMM, the approach should produce sensible outputs. A GMM is a reasonable model given that single cell data is known to often form clusters, and we have shown that the individual metrics (after appropriate transformations) typically follow Gaussian distributions. Non-Gaussian and non-parametric approaches to outlier detection could also be considered, such as estimating cell densities in QC space and then excluding cells in sparsely populated regions. However, this would risk excluding cells that were only rare in that sample, and our observations suggest that a GMM is reasonable for various experiments. By fitting one model across multiple samples, `SampleQC` is able to preserve celltypes that are abundant in some samples, but have low abundance in others and are therefore discarded by methods that operate only at the sample level.

Although the downstream effects of choice of QC metrics are not well understood, there are good biological motivations for the metrics currently in use. Low library sizes indicate droplets where the cell lysed before the reaction took place, droplets that contain only debris, or where the reaction was not successful, while high library sizes may correspond to multiple cells in one droplet. Number of features observed is highly correlated with library size, and therefore the same mechanisms for cell quality are relevant. Mitochondrial proportion may be high in cells that are stressed, or that lysed partially before measurement (resulting in a loss of cytoplasmic RNA and corresponding increased proportion of mitochondrial RNA); in single nuclei RNA-seq, the process of preparing nuclei should wash away mitochondria, meaning that mitochondrial reads may indicate contamination by ambient RNA. The relative proportion of spliced to unspliced reads indicates the relative abundances of nuclear and cytoplasmic RNA in a droplet (22). Unspliced reads should only be present in the nucleus; high ratios of spliced to unspliced read counts in single nuclei data therefore allow us to identify nuclei that were not successfully stripped of their cytoplasm, and are likely to be contaminated; meanwhile in single *cell* RNA-seq data, high proportions of unspliced (nuclear) RNA may indicate lysed cells. We expect that as new technologies are developed, new QC metrics will accompany them.

We have demonstrated `SampleQC` on standard measures of cell quality. However, the model is flexible, and can be applied to any set of metrics specified by the user that approximately follow a Gaussian mixture distribution. We also note transformations that allow non-Gaussian metrics to be transformed into spaces where they are more likely to be Gaussian. For example, `SampleQC` transforms mitochondrial proportions via the logit function, and spliced:unspliced ratios via log. `SampleQC` is effectively a general outlier detection technique for GMMs, and this should allow `SampleQC` to be applied to datasets with other or multiple modalities. We have applied `SampleQC` to CITEseq (Figure S10), but in principle, the framework can be applied to data from scATACseq (24), scCATseq (26), single cell techniques based on combined modalities, and spatial transcriptomics, by selecting appropriate measures (e.g., antibody tag depths for CITE-seq, sparseness for scATAC-seq, etc.). More exploration may be required to identify the set of QC measures that are sufficient to filter questionable cells, also because many existing measures are highly correlated, which could mean they are redundant.

QC does not need to be conducted entirely at the filtering stage. Another option is to carry forward more cells to a clustering/annotation step and then filter out subpopulations that are of debatable quality, assuming some biological knowledge can be used to assign this. And here, it is difficult to know if there is a clear “optimal” path. On the one hand, including more sparser cells may have adverse effects on similarity calculations, estimation of highly-variable genes, or on the low dimensional projections that clustering is based on; on the other hand, `SampleQC` summaries (e.g., P-values representing cells in low-density regions of their QC space) could be used afterwards as indicators for removal of individual clusters. The effect of including sparse cells on direct cell type annotation methods is not well studied. Furthermore, it is not clear at what point entire samples should be removed from downstream analysis; while `SampleQC` gives useful visualizations to characterize the QC of many samples, a judgement call is still needed.

It is worth noting that our performance results depend on the choices made in implementing the simulation, largely because we are not aware of other frameworks to simulate scRNA-seq QC summaries. Ultimately, our simulation highlights that inference from our model is possible, but does not directly show that cells are ‘bad’. While we tried to represent parameters and phenomena observed based on our experience in large scRNA-seq datasets, further development of the simulation framework would be desirable. We note that as the underlying models for both `scater` and `miQC` are unimodal, any circumstance where there are multiple distinct healthy cell populations may result in a performance advantage for `SampleQC`.

We do not provide any statistical guarantees that `SampleQC` converges to the optimal solution. However, `SampleQC` does provide automatically generated and comprehensive reports that show the fitted model over the empirical density. The models defined are conceptually simple, and the reports should therefore allow users to check both the validity of the fits and the appropriate number of clusters for each group.

We also do not attempt to solve the problem of selecting the appropriate value of *k*; this is a challenging and extensively-discussed problem in statistics (21). As noted above, the automatic reports allow users to inspect whether the model has fit appropriately. In addition, using robust estimators makes the procedure less sensitive to misspecification of the number of components: robust methods expect some of the data to be outliers, and therefore do not attempt to ‘explain’ all data; non-robust methods attempt to explain the whole dataset, and where the number of components is misspecified, the estimated means may lie between components.

As experiments increase in size and complexity, the task of removing unwanted cells becomes correspondingly more challenging, especially if users would like to preserve rarer populations. `SampleQC` attempts to solve this problem, by first identifying groups of samples with similar QC distributions, then fitting an empirically motivated statistical model to the QC metric data for each group. The model is simple, meaning that it is fast to fit and easy to check. This allows users to obtain an overview of data quality for their entire experiment, and to accurately identify cells for exclusion without having to make decisions regarding every sample individually. Our intention is to provide a fast, flexible and easy-to-use method that allows investigators to do quality control with minimal loss of their valuable data.

## Methods

### SampleQC model

`SampleQC` is based on standard approaches to identifying GMMs, with two important changes. Firstly, the model includes terms that are sample-specific: each sample has a sample shift term, i.e. a uniform change in values applying to all cells in a given sample. In addition, while we assume that each sample has the same mixture components (= QC celltypes), the *proportion* of these components may vary within each sample. For cell *i*, which comes from sample *j* in QC celltype *k*, this gives us the following model for the QC values *X*_*i*_:

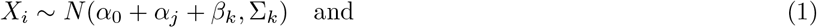

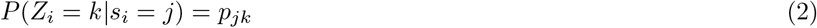

where

*s*_*i*_ = *j*, cell *i* is from sample *j*

*Z*_*i*_ = *k*, cell *i* is in mixture component *k*

*α*_*j*_ sample shift for sample *j*

*β*_*j*_ mean for mixture component *k*

Σ_*j*_ covariance matrix for mixture component *k*

*p*_*jk*_ mixing proportions in sample *j*

Secondly, the fit is robust, in the sense that it is less sensitive to outliers. The standard approach to fitting GMMs is via expectation-maximization, where the method alternates between fixing the means and covariance matrices then identifying the expected values for the latent cluster variables (‘expectation’), and fixing the latent cluster variables then calculating the maximum likelihood estimators (MLEs) for the means and covariances (‘maximization’). This approach assumes that all observations come from the GMM distribution. If the data is contaminated by a small proportion of observations that do not come from the assumed distribution (outliers), then this procedure may fail, especially regarding estimation of covariance matrices, in turn affecting cluster membership identification (19). To mitigate this problem, we use a robust method to estimate the means and covariance matrices, based on using the points nearest the centre of the component (20). Robust estimators are designed to return good results even in the presence of outliers, which is especially important when the principle assumption of our task is that outliers may be present.

In addition to identifying outlier cells, fitting the GMM gives in estimated statistics as outputs, namely: the QC metric shift for each sample 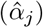, the location and shape of each mixture component 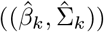, and the proportions of the components in each sample (*p*_*jk*_). These can be used to identify samples with markedly different statistics to others in their sample group, both in terms of mean QC values, and in terms of the split of cells across QC celltypes. These samples are therefore flagged to experimenters for detailed checking and potential exclusion.

### Criteria for outliers

In the software (see below), the user can also specify the threshold of the *χ*^2^ distribution to determine the how stringent the outlier exclusion should be. In particular, under the null of no outliers, the Mahalanobis distance (to the centre of the nearest GMM component) should follow a *χ*^2^ distribution (with degrees of freedom equal to the number of QC metrics used as input). Large distances are therefore by definition very unlikely under the GMM model and represent the candidates to filter out. This can be achieved by setting a threshold on the tail of the *χ*^2^ distribution

### Software

The input to `SampleQC` is a data.frame where each row corresponds to a cell; the columns must contain the chosen QC metrics and sample labels, and can optionally also include additional sample annotations such as batch or experimental condition. `Using a data.frame` as input makes the software accessible to users of both the `SingleCellExperiment` and `Seurat` packages (and, by saving intermediate files as CSV, to users of `scanpy`). Please see the user guide for further details.

Prior to running `SampleQC`, we recommend excluding empty droplets, for example by running emptyDrops with either the default or a more generous cutoff, to ensure that all genuine cells are included. This may result in some empty droplets being used as input to `SampleQC`, but they should be identified as outlier cells and later excluded.

We then recommend removing multiplets *before* running `SampleQC` for two principal reasons. Firstly, excluding low quality cells first assumes that all doublets are composed of two good quality cells; by excluding low quality cells first, it is then more difficult to detect doublets comprising one good quality and one low quality cell. Secondly, just as the count vectors for doublets are linear combinations of two true cells, their QC metric values are also functions of combinations of two true cells. The practical impact of this is that doublets introduce a small amount of noise to the QC metric distributions; excluding them results in cleaner distributions that are slightly easier for `SampleQC` to fit. (In addition, the number of cells used in the run is needed to accurately estimate the expected number of doublets. Excluding low quality cells first would distort this calculation; although this can be addressed by recording the initial number of cells.)

To run `SampleQC`, the user first runs the function `calculate_pairwise_mmds`, which estimates MMD values for each pair of samples, identifies groups of samples with similar QC metric distributions, and calculates embeddings in lower-dimensional spaces (both MDS and UMAP). The user can then plot the samples and check for evidence of batch effects, e.g. if all samples from a given well, run or experimental condition are grouped together. Users can also check whether any sample group has consistently low quality cells (e.g. high mitochondrial proportions) and consider excluding all samples in that group.

To run the cell outlier section of `SampleQC`, the user then selects `qc_names`, the QC metrics to use, and *k*, a vector corresponding to the number of mixture components for each sample group, and calls the function `fit_sampleqc`. `SampleQC` then fits a GMM to each sample group, and uses cell likelihoods to identify outlier cells. Users would typically start by using *k* = 1 for all sample groups; as the simplest model, this is extremely fast. The report includes biaxial plots of the QC metrics for all samples, which allow users to judge what value of *k* is most appropriate for each sample group, then rerun `SampleQC`. Users can inspect the cells identified as outliers by generating an html report with the function `make_sampleqc_report`. The user can also specify `alpha`, the *χ*^2^ distribution threshold determining the how stringent outlier exclusion should be, however this is less critical and the default value (`alpha` = 0.01) is generally appropriate. alpha specifies the distance away from the centre of the nearest GMM component that should be considered as an outlier. Under a robust model, the distribution is assumed to be a GMM plus a contamination component that does not follow the GMM, and is therefore by definition very unlikely under the GMM distribution, so as long as `alpha` is set reasonably low (e.g. 1%, 0.1%), this will only have marginal effects on results.

If a large proportion of a sample is of poor quality, `SampleQC` may identify these cells as one sample cluster, and one of the mixture model components may therefore be centred on them (e.g. the cells with high splice ratio in Figure 5). Without intervention, these would be reported as good cells rather than outliers. `SampleQC` therefore includes functionality allowing users to specify that any component whose mean meets a certain criteria should be regarded as outliers. For example, a component whose mean mitochondrial percentage is greater than a given percentage can be labelled as outliers.

### Simulations

We developed a framework for simulating samples of QC data that reflects the complexity observed in real datasets, following the model given in Equation 1, but additionally modelling outliers; in the `SampleQC` R package, users can access this functionality from the `simulate_qcs` function. For the simulations, we assume that there are three QC metrics: log library sizes, log feature counts and `logit` mitochondrial proportion. The simulations are hierarchical: firstly, parameters for group size and GMM components are drawn at the whole-experiment level; then, we draw parameters for sample quality at the sample group level, capturing variability that is coherent across groups of samples; then we draw parameters for each individual sample and QC metrics for each individual cell. To model the QC metrics, we assume that most cells are healthy, i.e. they derive from the experiment-level GMM, while a small number are ‘bad’, i.e. their values are perturbed away from the means of the GMM. These steps are set out in Algorithm 1.

At the experiment level, we simulate the covariance matrices and relative means of the mixture components (QC celltypes) that are present in the experiment (Algorithm 2). We then simulate parameters reflecting coherent behaviour across different groups of samples. First, we partition the cells between the groups according to a multinomial distribution, simulate how many samples are present in each group, and randomly assign QC celltypes to each group. We assign the QC celltypes such that each group has at least one QC celltype present, and that no two groups have identical sets of QC celltypes. To simulate coherent differences in outliers between each group, we simulate parameters from a beta distribution that determines how the proportion of outliers varies across samples (`p_out_0, theta_0/`), and how perturbed these outliers will be relative to healthy cells (`p_loss_0`, the mean proportion of non-mitochondrial reads that is lost).

We then simulate samples and cells within each sample group (Algorithm 4). Within each sample, we first simulate the sample shift from a multivariate Gaussian, and simulate the proportions of the different QC celltypes via a Dirichlet distribution (`p_jk`). We also simulate what proportion of cells in this sample are outliers, `p_out_j`, from a beta distribution with parameters (`p_loss_0, theta_0`), and the extent of perturbation in unhealthy cells, `p_loss_j`, from a logit-normal distribution.

We can then simulate each healthy cell, by first drawing its QC celltype from a multinomial with **p** = {`p_jk`}, then drawing from multivariate normal (`beta_k, Sigma_k`) for the appropriate *k*. Adding this to the relevant mu_0 gives us the QC metrics for a healthy cell. We then decide whether this cell is healthy or not by drawing from a binomial distribution (with probability `p_out_j`. If the cell is healthy, its QC metrics are left unchanged. If not, its non-mitochondrial reads are randomly decreased by a proportion `p_loss_j`. This results in both a reduction in total counts, and an increase in mitochondrial proportion. We apply the same random reduction to the number of features.

The outputs from the simulations show similar distributions of QC metrics to real data (Figure 5).

#### Algorithm 1: simulate_qcs: Overview of simulations (* indicates variables that would be known and used as input for real data)

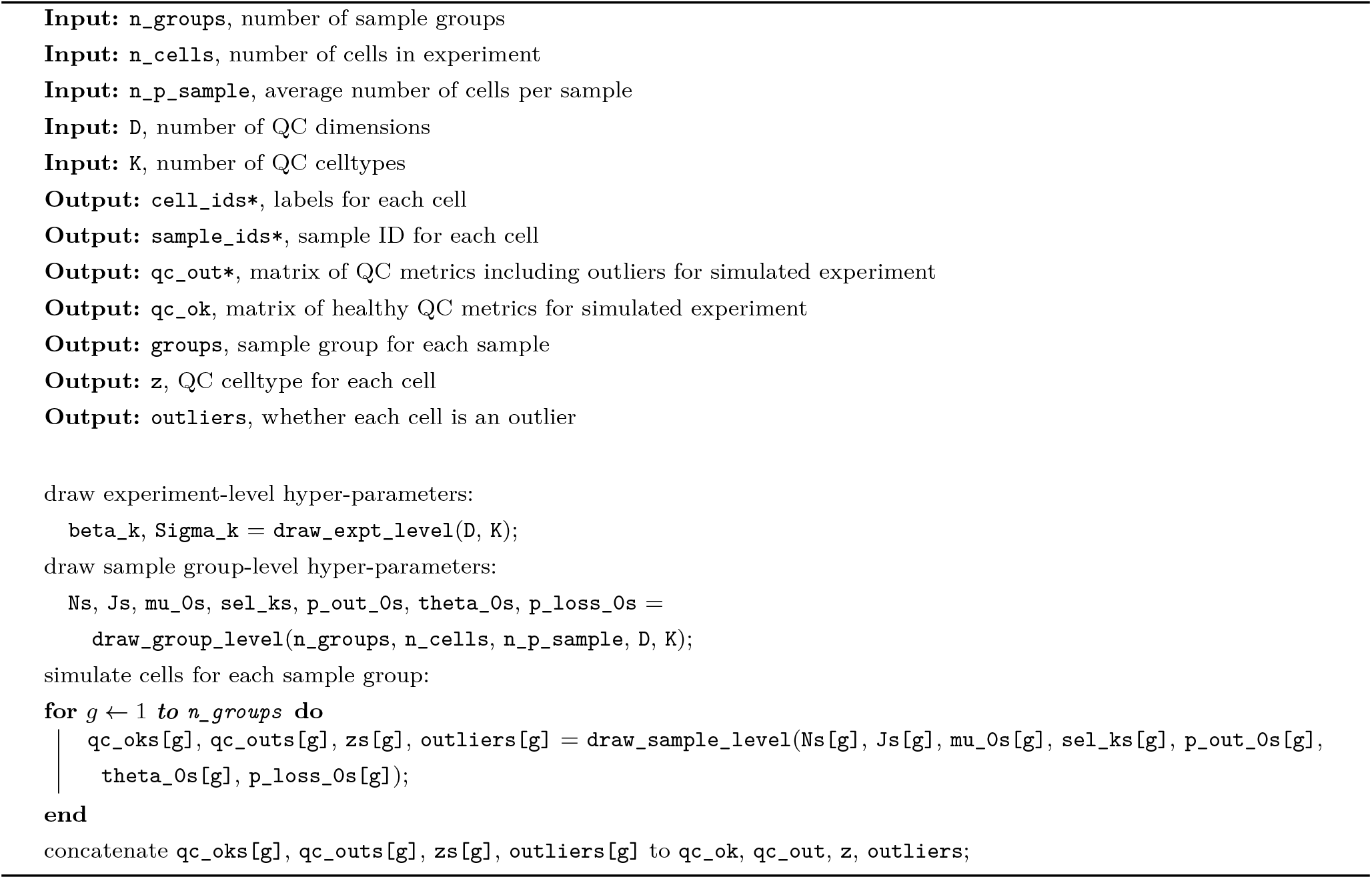

#### Algorithm 2: draw_expt_level: Pseudocode for drawing experiment-level parameters

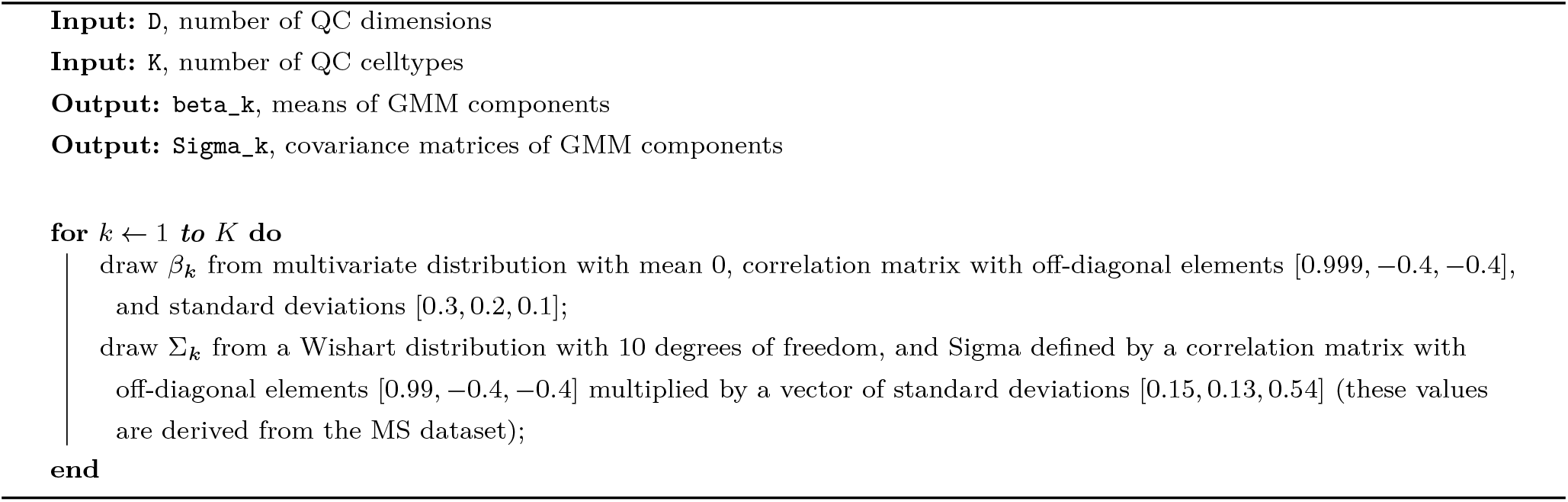

#### Algorithm 3: draw_group_level: Pseudocode for drawing sample group-level parameters

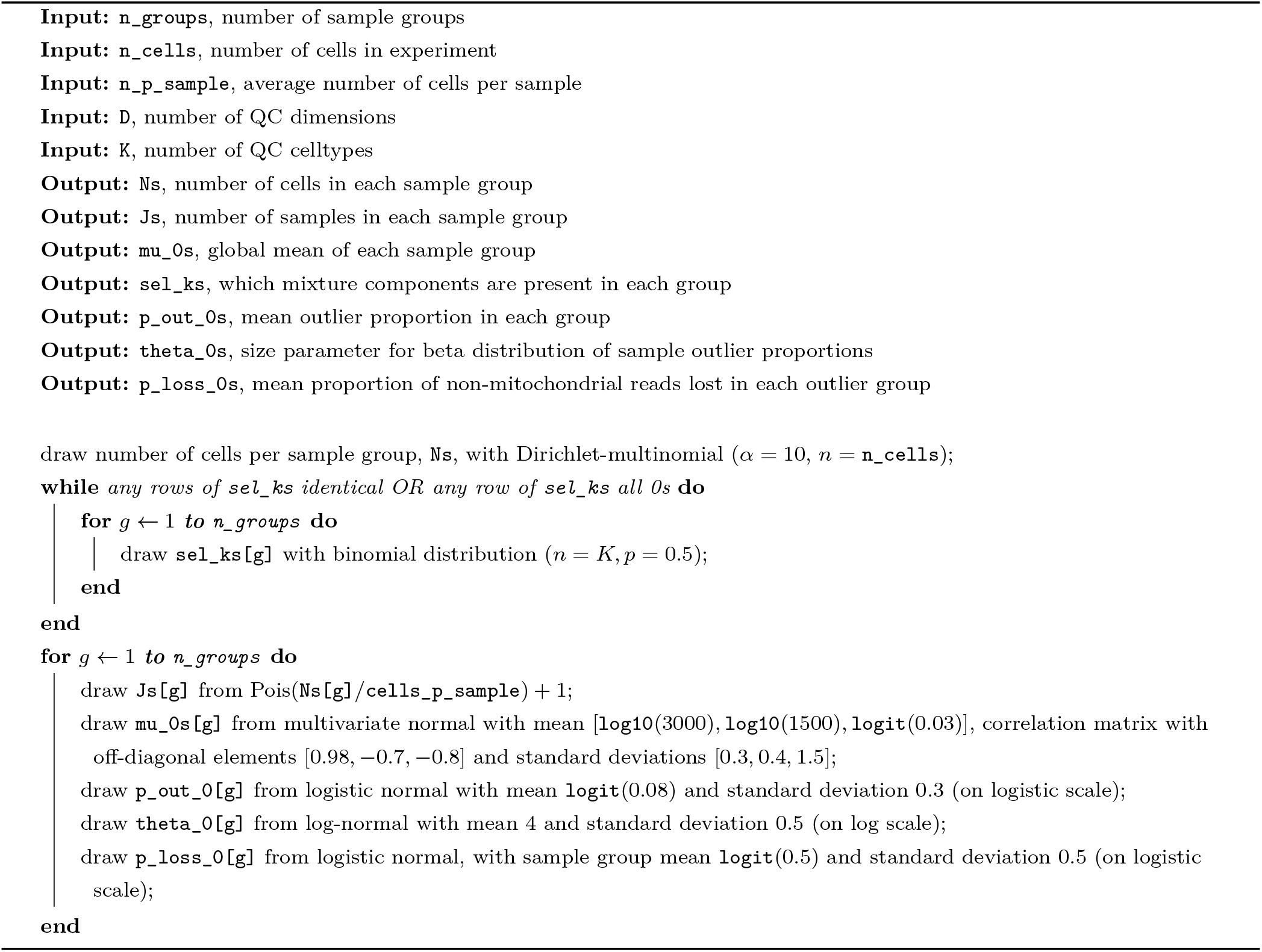

#### Algorithm 4: draw_cell_level: Pseudocode for drawing cell-level parameters

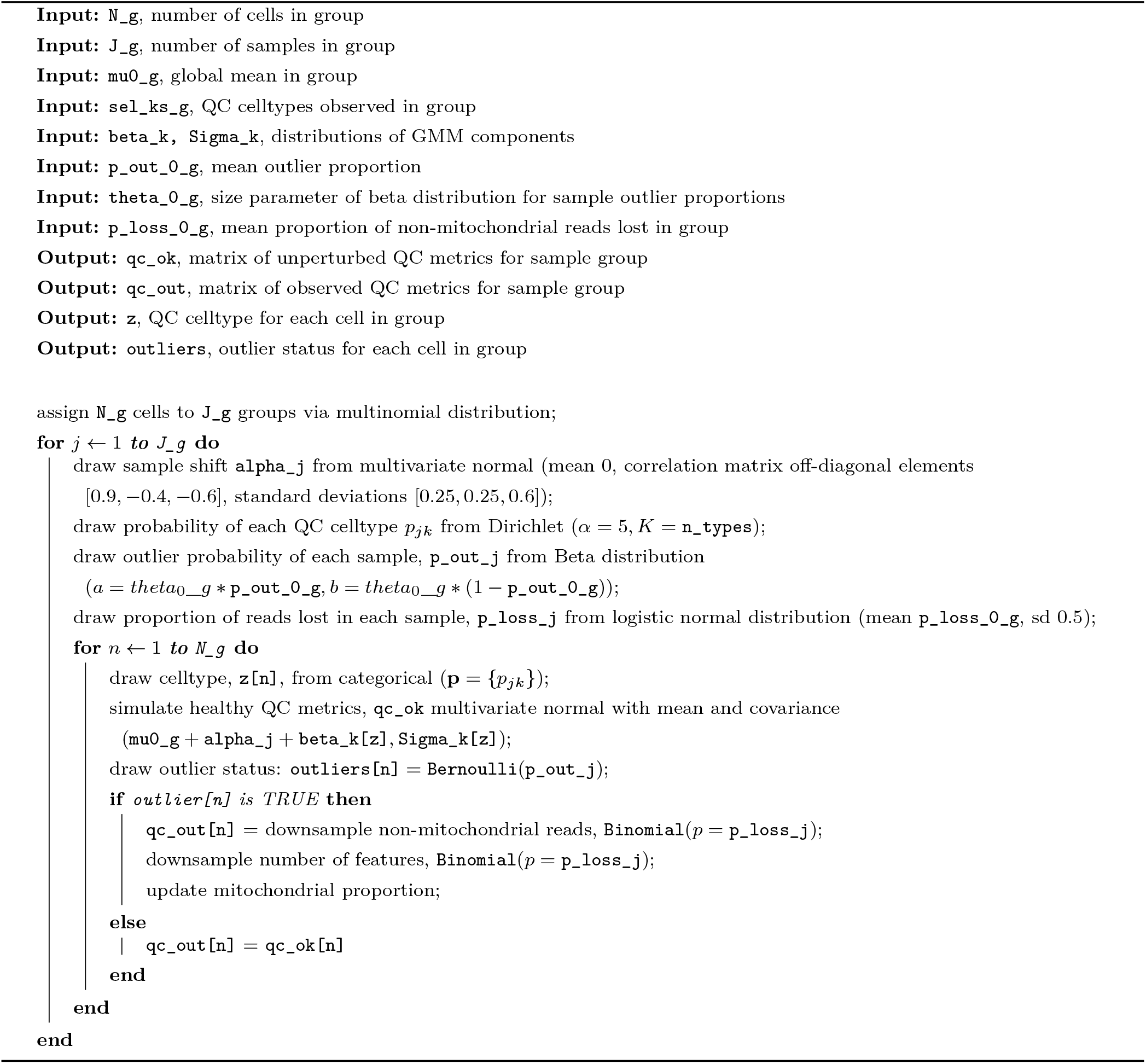

## Datasets

The datasets listed in Table were used in this paper, obtained via the `scRNAseq` package (27).

## Author’s contributions

We use the CRediT taxonomy to define author contributions:

- WM: Conceptualization, Methodology, Software, Formal analysis, Investigation, Visualization, Data curation, Writing - Original Draft, Writing - Review & Editing
- MDR: Supervision, Methodology, Writing - Review & Editing, Funding acquisition

## Acknowledgements

We thank Helena Crowell, Pierre-Luc Germain and Simone Tiberi for their valuable feedback on the manuscript and software. We thank Dheeraj Malhotra and Roche for the large experimental dataset (see Data availability below).

## Data availability

The data in Figures Figure 4 and Figure 5 is QC data taken from human brain tissue, including samples from patients with neurodegenerative conditions. The data derives from a study undertaken as part of a research collaboration agreement between the University of Zürich and F. Hoffman La Roche AG. The study is not yet published and the experimental labels have therefore been anonymized. Although we have made the anonymized QC metric data available (see DOI: 10.5281/zenodo.5258847), the full expression data is not yet public.

## Code availability

The `SampleQC` software package is accessible at https://github.com/wmacnair/SampleQC; a code snapshot of the software package is available here: DOI: 10.5281/zenodo.6414310). A repo for reproducing the analyses in the paper is accessible at https://github.com/wmacnair/SampleQC_paper_analyses and the corresponding HTML reports are available from https://htmlpreview.github.io/?https://github.com/wmacnair/SampleQC_paper_analyses/blob/main/docs/index.html; a code snapshot of the analysis repository at time of submission is available here: (see DOI: 10.5281/zenodo.6414318)

## Tables

**Table 1:** List of datasets used in paper. All datasets are single cell RNA-seq unless otherwise stated.

| Short name | Tissue | Reference |
| --- | --- | --- |
| Campbell | Mouse brain | (28) |
| CITEseq | Human PBMC (CITE-seq) | (25) |
| HGSOC | Human ovarian cancer | (29) |
| Macosko | Mouse retina | (2) |
| Roche | Human brain (single nuclei RNA-seq) | (30) |
| Shekhar | Mouse retina | (30) |
| Wang | Human endometrium | (31) |
| Zeisel | Mouse brain | (32) |

## Supplementary Figures

**Figure S1:**
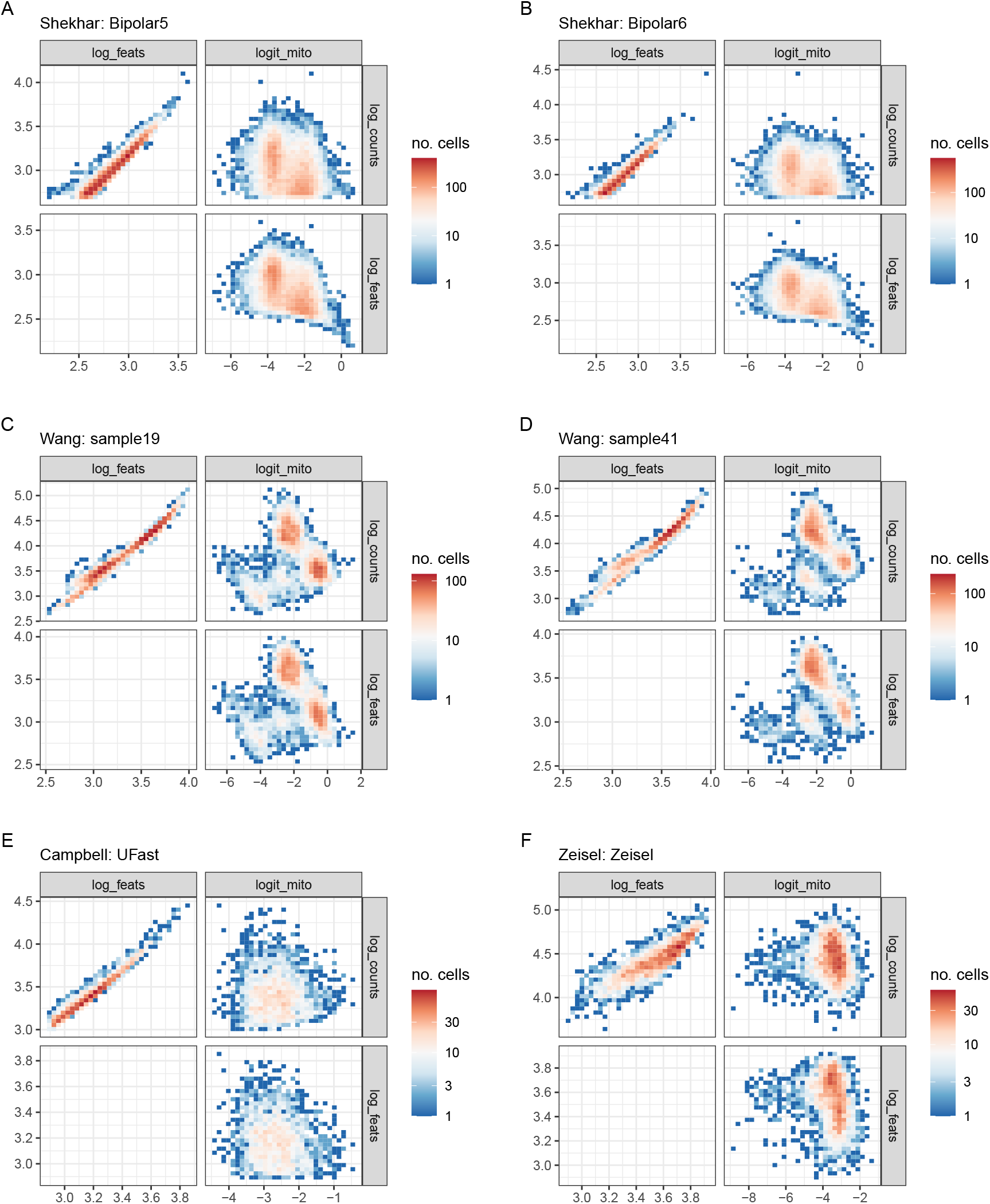
Biaxial empirical density plots of QC metrics from selected samples and datasets. Colour indicates concentration of cells with the same QC metrics. *log_counts* indicates log10 of library sizes, *log_feats* indicates log10 of number of detected genes, *logit_mito* indicates qlogis i.e. inverse logistic function of mitochondrial proportion. For list of datasets, see Table.

**Figure S2:**
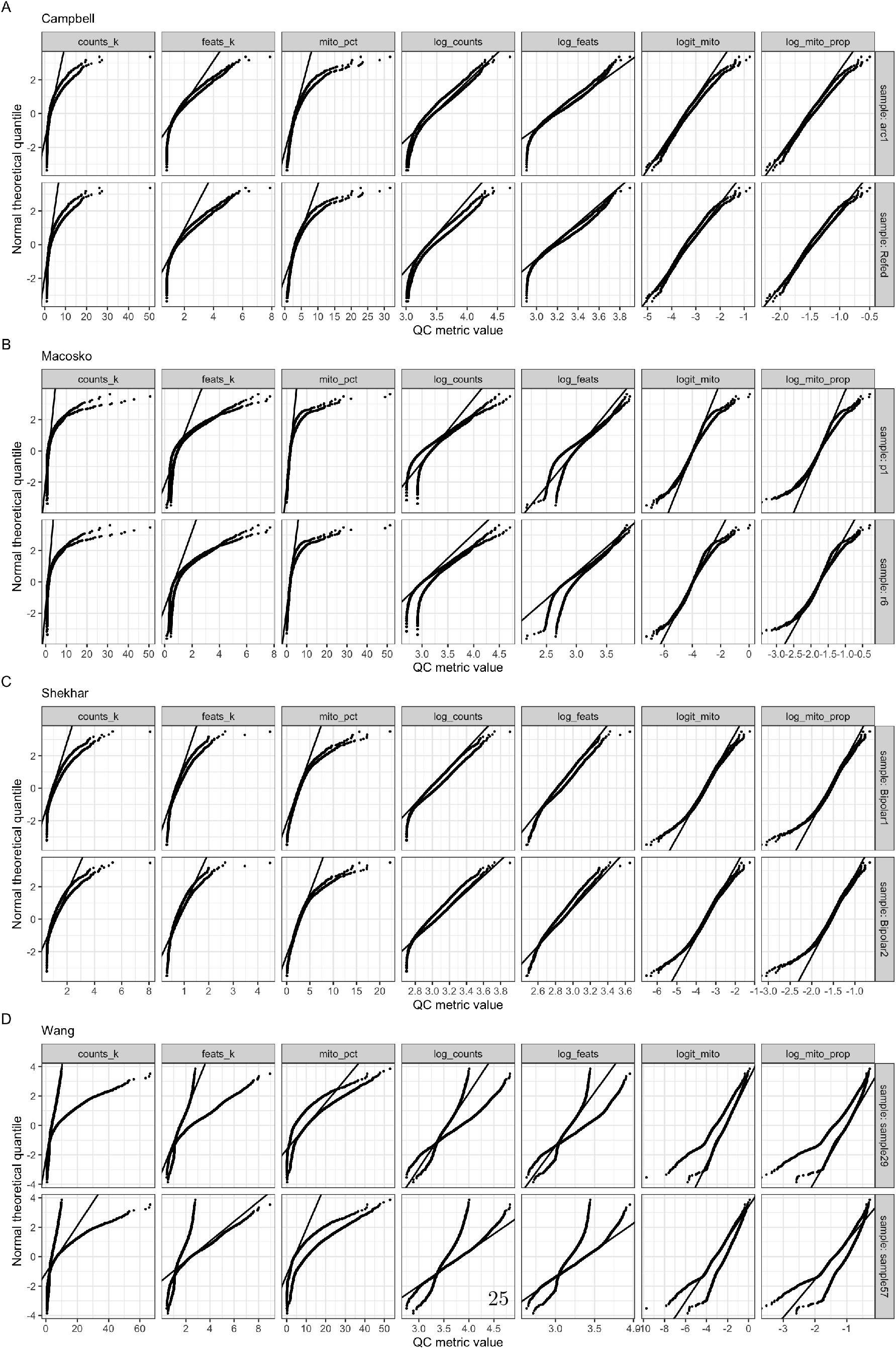
QQ plots of QC metrics from selected samples and datasets. y-axis is theoretical quantiles of normal distribution, using mean and standard deviation estimated from transformed QC metrics. Median of QC metric used to estimate mean, MAD of QC metric for standard deviation. x-axis is observed values of selected QC metrics in each sample. *counts_k* indicates library sizes in thousands, *feats_k* indicates number of detected genes in thousands, *mito_pct* indicates percent of mitochondrial reads, *log_counts* indicates log10 of total reads per cell, *log_feats* indicates log10 of number of detected genes, *logit_mito* indicates qlogis i.e. inverse logistic function of mitochondrial proportion, *log_mito_prop* indicates log10 of proportion of mitochondrial reads. For list of datasets, see Table.

**Figure S3:**
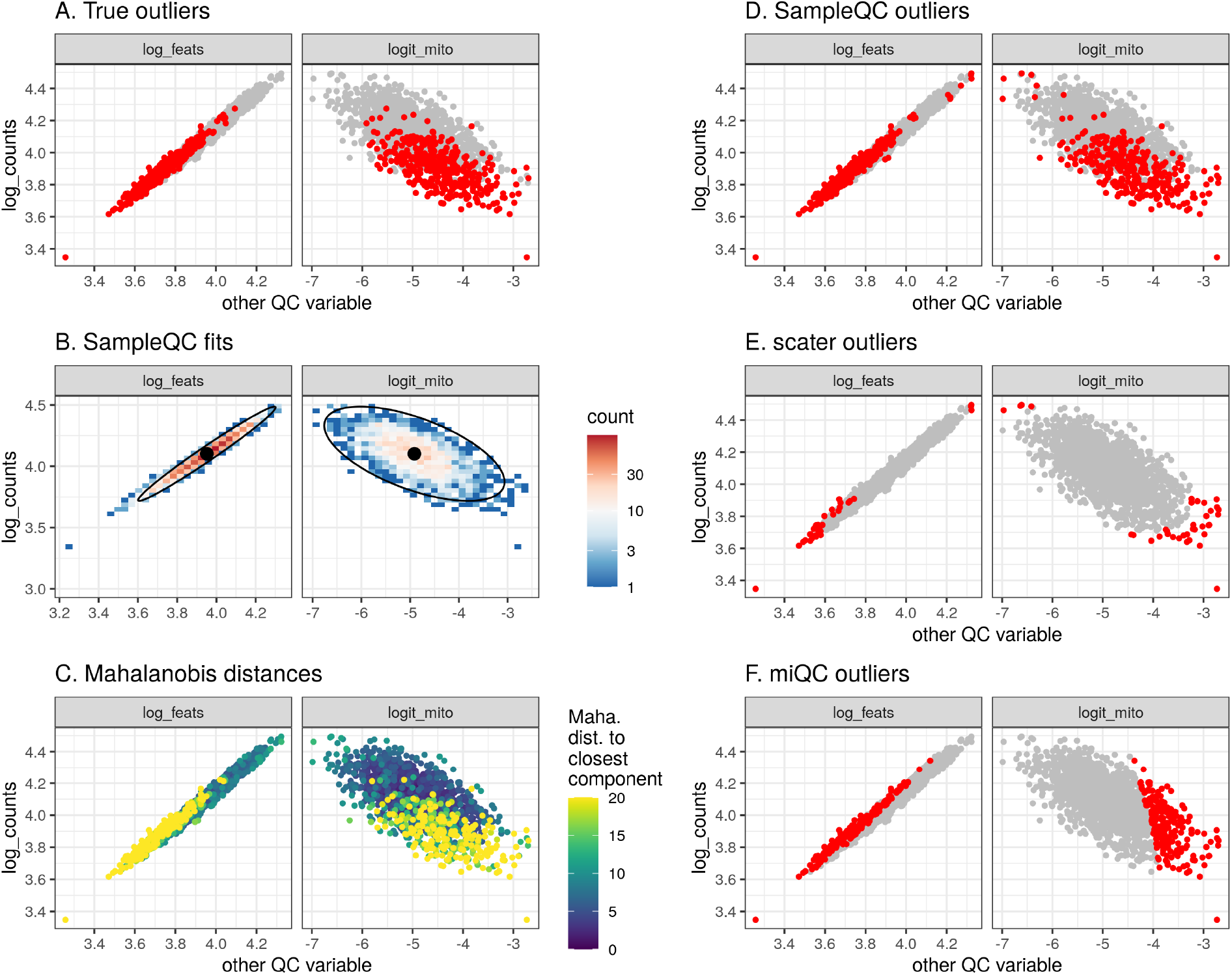
QC method comparisons on unimodal simulated data. **A** True simulated outliers (red). **B** Multivariate density plot of simulated data, showing `SampleQC` fits. **C** Mahalanobis distance to nearest cluster under fitted `SampleQC` model. **D, E, F** Outliers detected by `SampleQC`, `scater` and `miQC` respectively.

**Figure S4:**
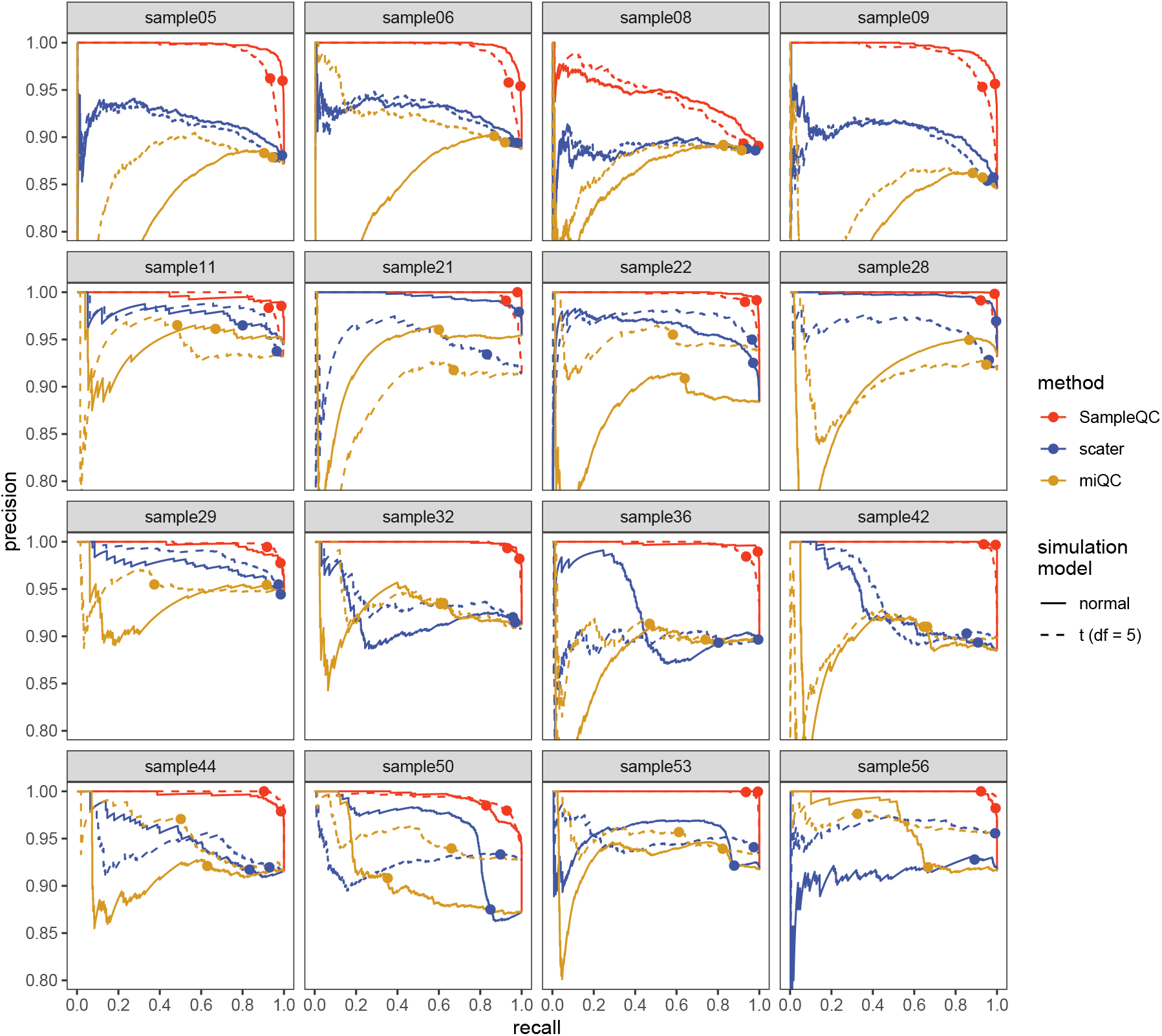
PR curves for individual samples. PR curves for 20 randomly selected samples from simulated experiments. Precision is proportion of cells reported as ‘good’ that are actually ‘good’; recall is proportion of all ‘good’ cells that are reported as ‘good’. Samples in first row are unimodal, i.e. they are composed of one celltype; other samples have either 2 or 3 celltypes. Dashed lines show results for simulations where QC celltypes have QC metrics modelled as multivariate t-distributions with 5 degrees of freedom, and other simulation parameters were kept constant.

**Figure S5:**
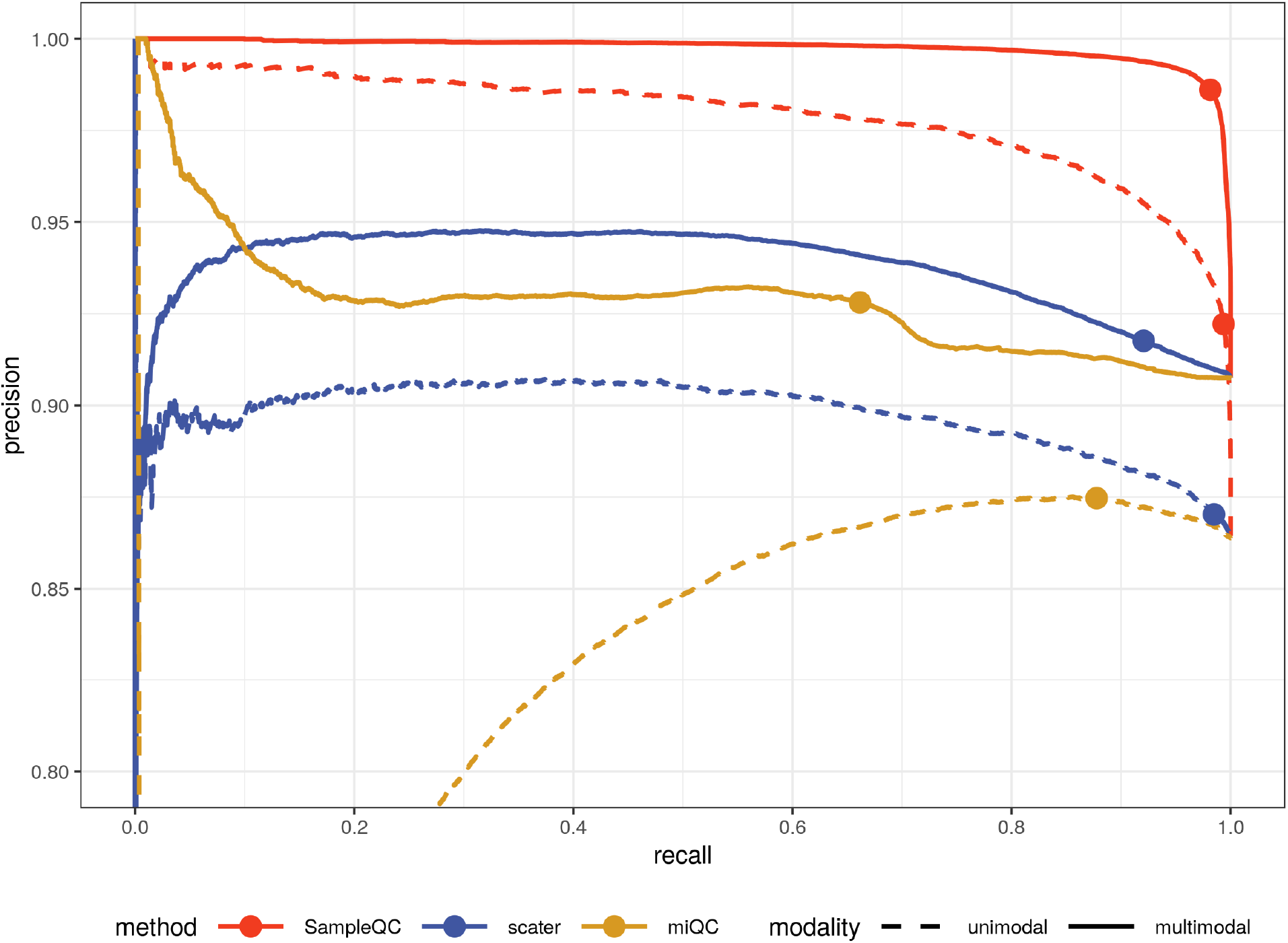
Precision-recall curves split by number of celltypes present. Modality indicates number of modes present in the distribution, i.e. the number of celltypes.

**Figure S6:**
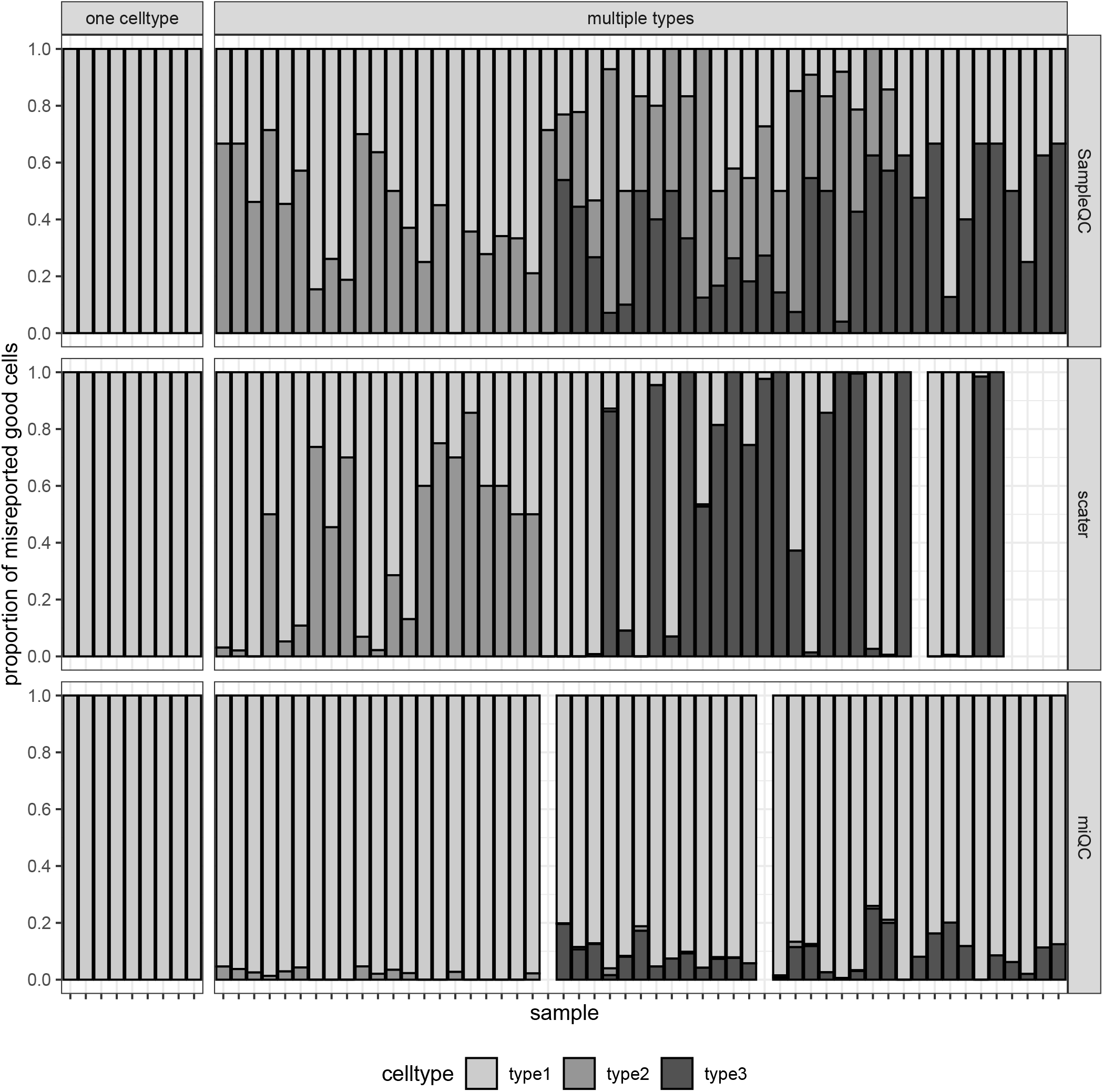
Split of misreported good cells by celltype.

**Figure S7:**
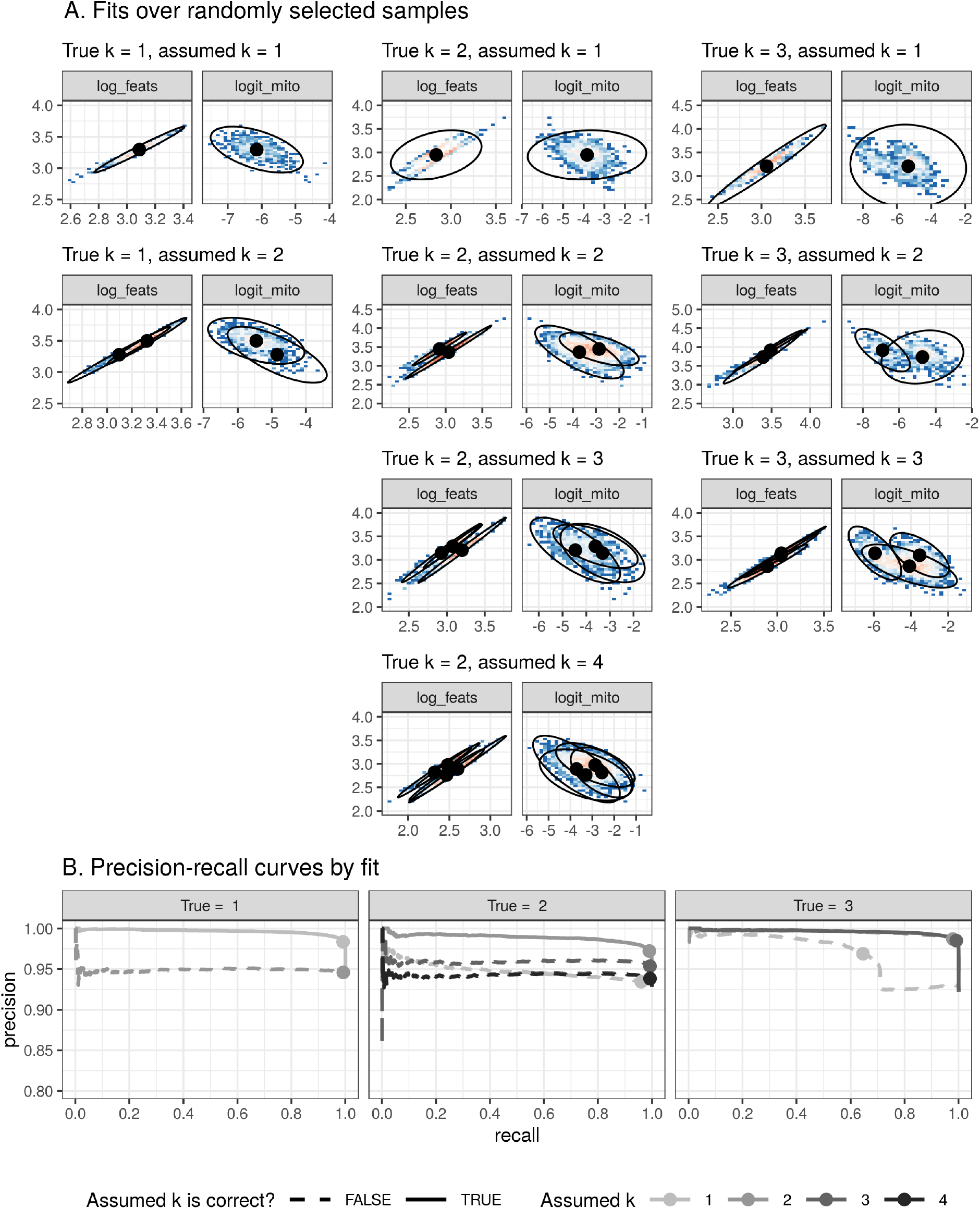
Robustness to choice of *k*. **A** `SampleQC` fits to simulated data, for combinations of `k_fit` and `k_sim`; columns are `k_fit`, rows are `k_sim`. Each fit illustrated by randomly selected sample for each combination. Note that for some combinations of `k_fit` and `k_sim`, `SampleQC` was not able to fit the data; these are blank. **B** Precision-recall curves of identification of ‘good’ cells. Solid lines show curve where `k_fit` = `k_sim`. Dots show default cutoff (*<* 1% Chi-squared tail).

**Figure S8:**
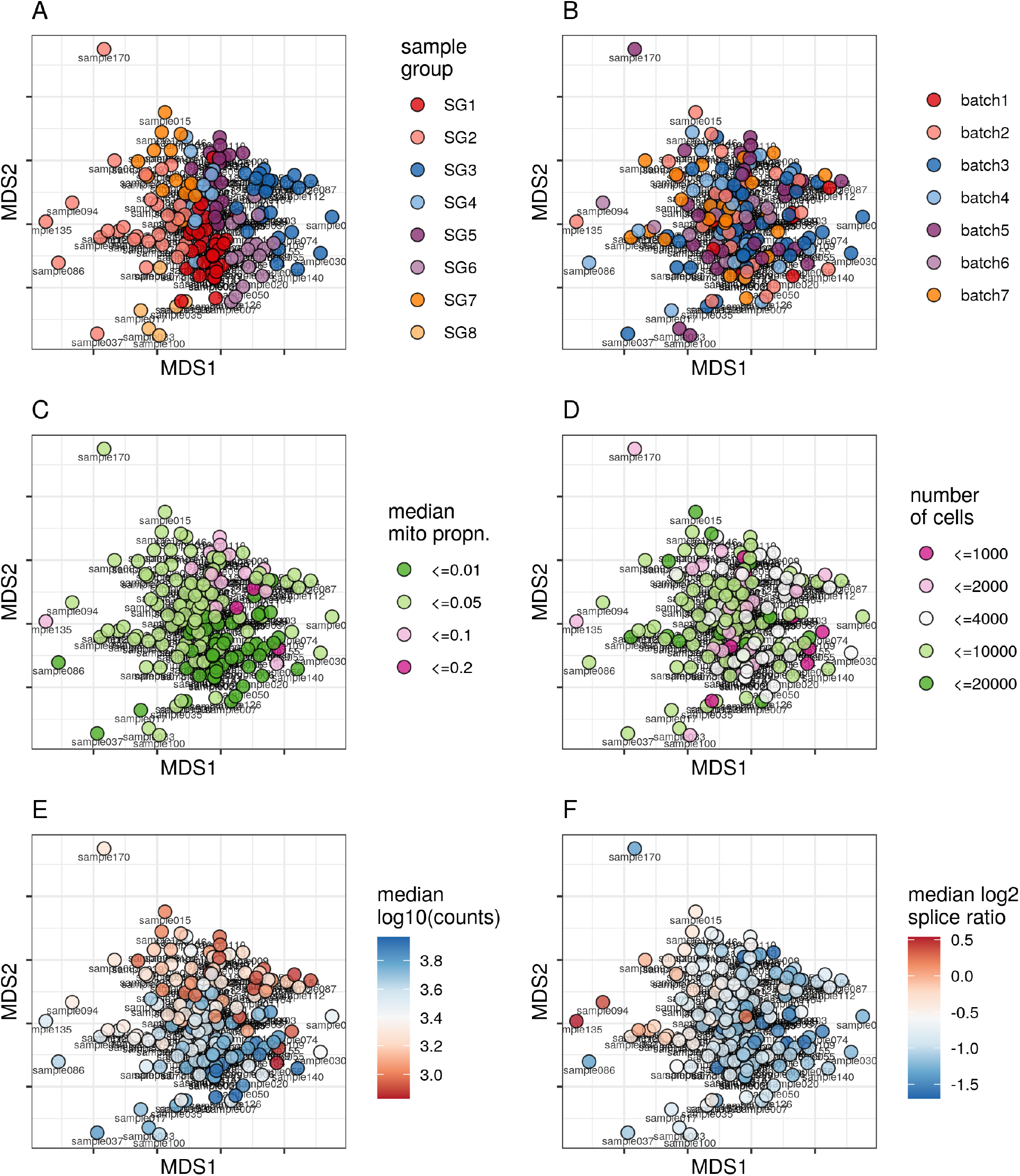
Sample-to-sample distance embedding (MDS). **A** Sample group, as derived from `SampleQC` clustering **B** Sequencing batch **C** Mitochondrial proportion (categorized) **D** Number of cells (categorized). **E** Median counts per cell **F** Median log2 splice ratio per cell.

**Figure S9:**
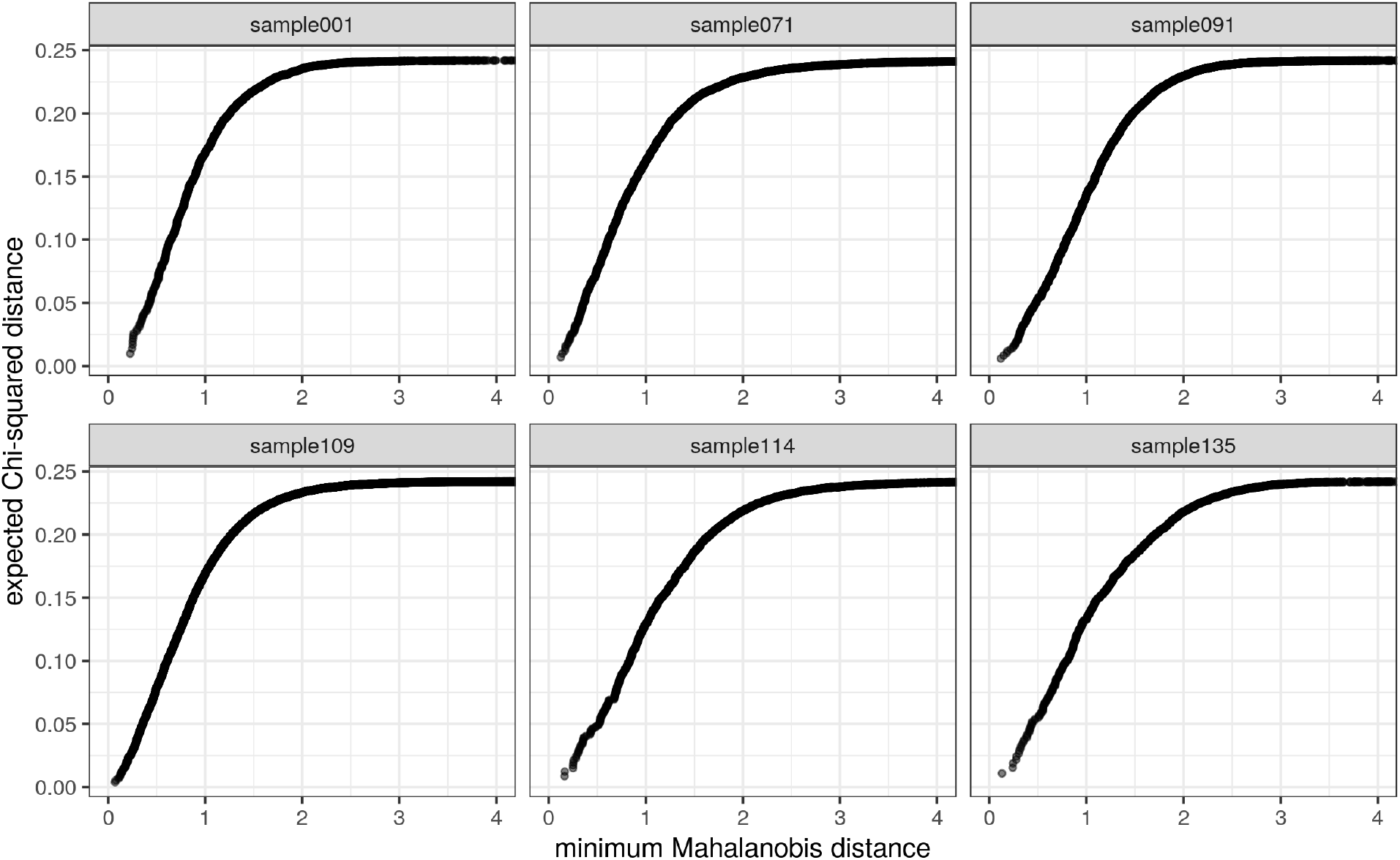
Comparison of Mahalanobis distances under `SampleQC` model and distances under expected Chi-squared distribution. Mahalanobis distances for Roche data and model fit in Figure 5, samples selected at random.

**Figure S10:**
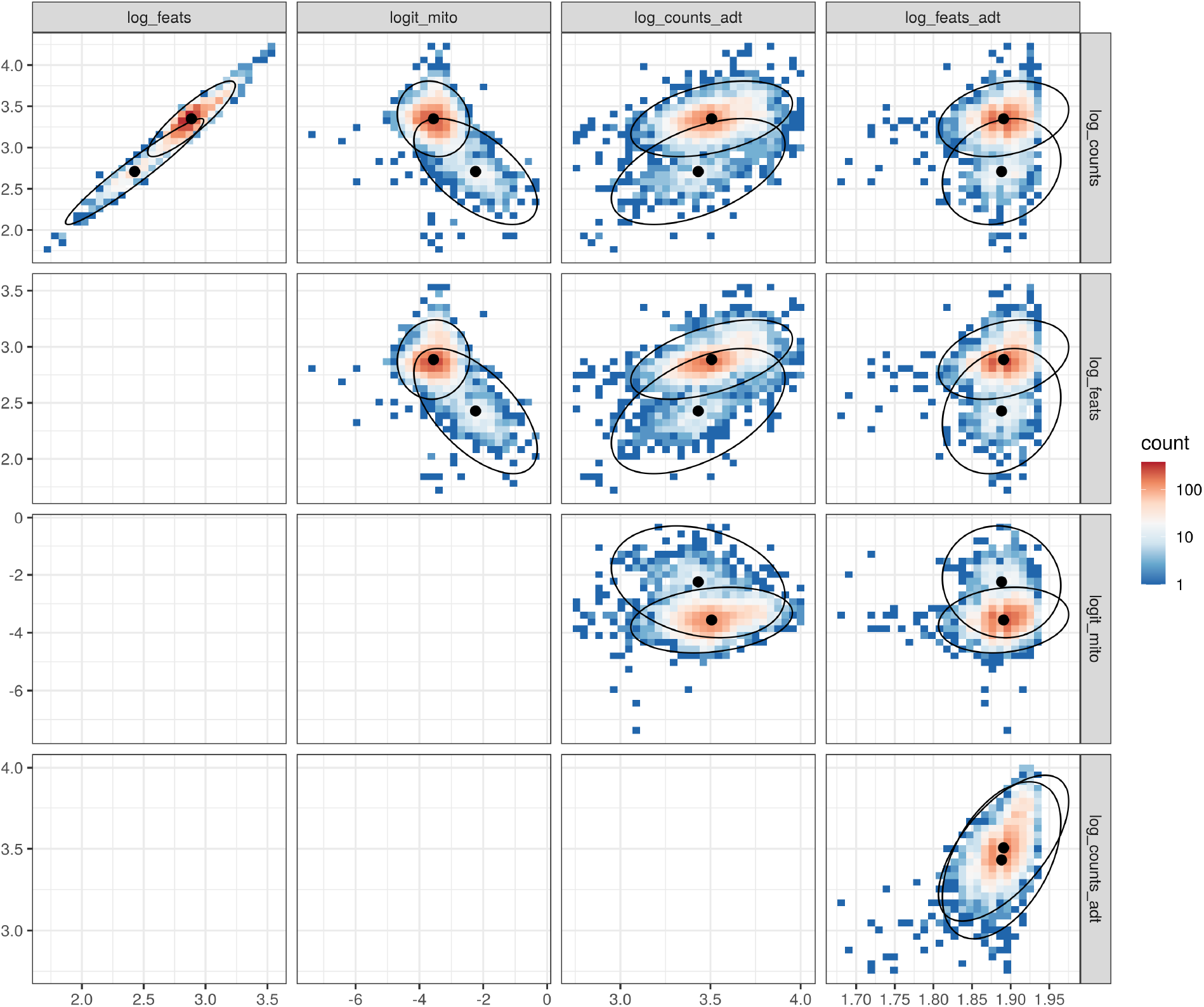
`SampleQC` outputs for selected CITE-seq sample. Biaxial distributions of CITE-seq QC metrics for PBMCs (25), annotated with fitted models fitted by `SampleQC`. Selected sample *H1B2ln2* from batch 2; distributions and fits for other samples are similar.

**Figure S11:**
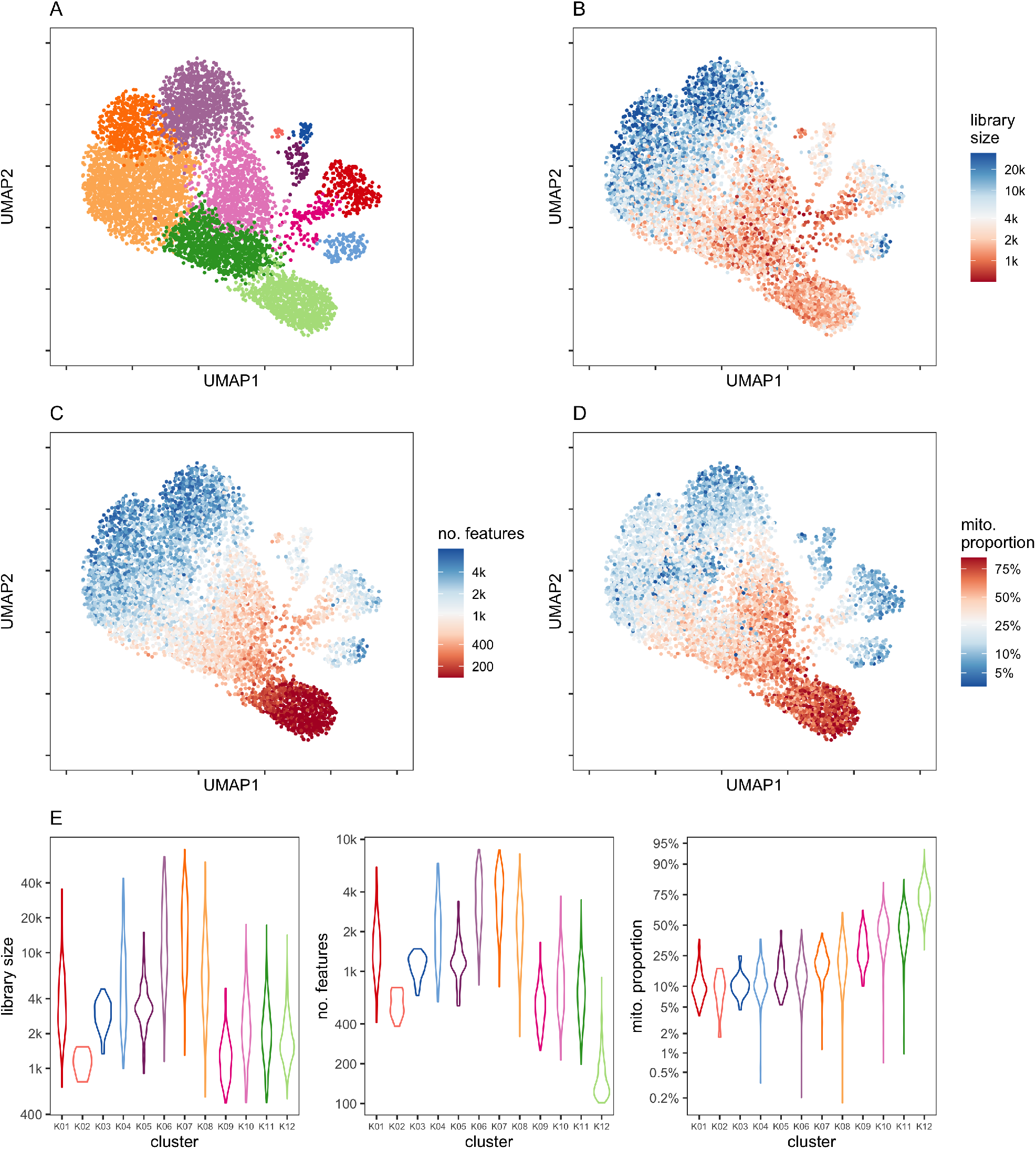
Identifying outlier cells via standard single cell clustering. Selected sample is ovarian cancer tissue, *16030×4* from the HGSOC dataset (see section). **A** UMAP applied to highly variable genes, coloured by Louvain clustering (16). **B-D** QC metrics for each cell, truncated to median + / - 2 MADs. **E** QC metric distributions of clusters in **A**.

